# Food Ration Affects mRNA Processing, Translation, Proteostasis, and Cytoskeletal Responses During Heat Shock in *Mytilus californianus*

**DOI:** 10.64898/2026.07.12.738034

**Authors:** L. Tomanek, R. F. Fabela, M. A. May

## Abstract

Despite a likely role in setting stress tolerance limits, food ration has received limited attention as an ecological factor affecting the cellular stress response (CSR). To study the interactive effects of food and temperature on the proteomic heat shock response, we acclimated intertidal mussels (*Mytilus californianus*) to four combinations of nearshore (low) and aquaculture (high) phytoplankton levels, combined with low (20 °C) and high (30 °C) aerial temperatures during daytime low tides. Mussels were then exposed to an acute (6 h), aerial heat stress (33 °C) and allowed to recover for 1 h and 25 h in pre-exposure conditions. Proteomic changes in the gill before and after heat shock were measured using label-free liquid-chromatography-mass spectrometry. Compared to other acclimation treatments, low-food-low-temperature (LTLF) mussels modified more splicing factors, heterogeneous nuclear ribonucleoproteins, ribosomal proteins, and translation initiation and elongation factors, suggesting systemic changes to RNA processing, selection, and translation. LTLF mussels also increased chaperones of the actin- and microtubule-associated cytoskeleton and proteins that mature along the endoplasmic reticulum to Golgi secretory pathway. Simultaneously, actin stress fiber formation at focal adhesions and the extracellular matrix, along with anchoring of microtubule-associated cilia, indicate a possible system-wide mechanical breakdown of the cytoskeleton. Signaling proteins causing cytoskeletal changes varied mainly in LTLF mussels and suggest a food-dependent role for prostaglandin synthesis. Overall, the acclimation-dependent proteomic changes show how thermal conditioning and food ration together will shape the CSR of mussels during heat waves.

**SUMMARY STATEMENT:** Variation in mRNA processing, translation, chaperoning proteins and stabilization of the actin- and microtubule-associated cytoskeleton establish food ration as a major environmental variable of the cellular stress response in mussel gill.

## INTRODUCTION

The cellular stress response (CSR) elicits a transcriptional and translational upregulation to cope with unfolding proteins, assembling a proteome that can cope with the proteostasis challenge (Hartl et al., 2011; Liu & Qian, 2014; Rosenzweig et al., 2019). While studies of the ecological role and evolutionary variation of the CSR have identified environmental drivers of cellular processes (Feder & Hofmann, 1999; Kültz, 2005) and shown that acclimation modifies the CSR (Somero et al., 2017; Tomanek, 2010; Tomanek & Somero, 1999), studies on food ration, especially with respect to thermal conditioning, have been limited. For example, while there is evidence that both mRNA and rRNA processing (including alternative splicing) are sensitive to heat stress and that food ration can reprogram RNA processing in yeast and human cell lines (Mushegian, 2017; Yost & Lindquist, 1986), the simultaneous effect of both has not been tested. Likewise, while it is established that molecular chaperones, including the chaperonin-containing TCP-1 (CCT/TRiC) chaperone complex, are essential for folding and protecting cytoskeletal proteins during acute stress, the simultaneous effect with food ration has not been investigated (Kuzu et al., 2025). And while it is established that elements of the proteolytic machinery, especially the ubiquitin-proteolytic system (UPS), are upregulated following heat stress in marine bivalves (Hofmann & Somero, 1995; Hofmann & Somero, 1996), the effect of food ration on the various elements of the UPS is not known. This is also the case for the actin- and tubulin-associated cytoskeleton in mussels, which changes under chronic and acute thermal stress (Fields et al., 2012; Tomanek & Zuzow, 2010). However, studies on caloric restriction and its effect on longevity have been associated with improved proteostasis of the endoplasmic reticulum in *C. elegans* (Matai et al., 2019) and the activation of the heat-shock transcription factor 1 in yeast (Choi et al., 2013), suggesting that food ration may modulate aspects of the CSR. And while the effect of food ration may be inferred indirectly through a change in metabolism on ATP-dependent cellular processes, or calorie-dependent processes, like those involving NAD-dependent deacylases (i.e., sirtuin), on the stress response in mussels (May & Tomanek, 2024; Vasquez et al., 2017; Vasquez et al., 2020; Vasquez & Tomanek, 2019), it has not been directly tested at a system level. While the answers are likely species—and even population specific, considering ranges of food availability—a food-dependent proteomic signature will provide novel insights into the plasticity of the cellular stress proteome. We address these questions by evaluating the proteomic response of intertidal mussels that have been acclimated to four food and thermal regimes during recovery from an acute heat shock.

The marine mussels, *Mytilus* spp., have been used as models in environmental stress response studies for decades (Bayne, 2017; Gosling, 1992). They are dominant members of rocky intertidal communities, where they act as a “foundation species”, creating microhabitats and protection for other organisms within the dense mussel beds (Lafferty & Suchanek, 2016). Earlier studies focusing on the abundance of heat shock proteins to acute or chronic stress (Anestis et al., 2007; Buckley et al., 2001; Hofmann & Somero, 1995; Hofmann & Somero, 1996; Ioannou et al., 2009; Roberts et al., 1997) have been expanded to a system level understanding of the stress response by transcriptomic, proteomic, and metabolomic studies in *Mytilus* (Apraiz et al., 2006; Connor & Gracey, 2012; Fields et al., 2012; Georgoulis et al., 2023; Gleason et al., 2023; Gracey et al., 2008; Place et al., 2011; Tomanek & Zuzow, 2010; Tomanek et al., 2012; Vasquez et al., 2017; Vasquez et al., 2020). These studies demonstrated the broad range of the CSR complement, but none investigated the effect of food or its interaction with thermal conditions despite evidence of differences in several biochemical indicators along intertidal or geographical food gradients (Dahlhoff & Menge, 1996; Dowd et al., 2013; Lesser et al., 2010; Petes et al., 2008; Phillips, 2004). In addition, laboratory studies showed that higher food ration can enhance thermal stress tolerance and feeding rates almost as much as acclimation to warmer temperatures in response to acute aerial heat stress (Fitzgerald-Dehoog et al., 2012; May et al., 2021; Schneider et al., 2010), but the cellular mechanisms behind this are understudied (May & Tomanek, 2024). Finally, documented changes in primary production in the North Atlantic and predicted changes in phytoplankton distribution in the California Current suggest an important role for food ration in the response of *Mytilus* to global change (Dai et al., 2023; Moore et al., 2018).

Here we present the proteomic response to acute heat stress, using a liquid-chromatography tandem mass spectrometry (LC-MS/MS), label-free quantification approach (Michalski et al., 2011). Our results present a strong effect of food-by-temperature acclimation on mRNA selection and splicing, ribosomal proteins affecting translation, a range of molecular chaperones folding cytoskeletal proteins and proteins that help mature proteins along the endoplasmic reticulum (ER)—Golgi secretory pathway, and protein degradation pathways. They also provide evidence for food-dependent dynamics of the actin- and microtubule-associated cytoskeleton and its connection to the extracellular matrix (ECM) as the main cellular protein structures being destabilized during heat stress. These changes may even lead to the immediate remodeling of stress-induced signaling pathways, specifically prostaglandin synthesis. Two companion papers cover the food-dependent changes in core energy metabolism, redox balance, chromatin modifications and DNA repair and how they are mediated by one-carbon metabolism (Fabela et al., in review) and how these changes are affected by food-dependent sirtuins and their inhibitors (May et al., in review).

## MATERIALS AND METHODS

### Experimental Design

The experimental design and animal husbandry have been previously described in May et al. (2021) and Fabela et al. (in review). We collected adult *Mytilus californianu*s (Conrad) from Montaña del Oro State Park, Los Osos, CA, USA, and acclimated the mussels at the California Polytechnic State University Center for Coastal Marine Studies (Avila, CA, USA) (Fig. 1A). Mussels were held in flow-through, ‘tidal-simulator’ tanks that permitted control of tide height (2 x 6-h low:high tidal cycles), feeding, 12-h light:dark circadian cycle, and temperature throughout the acclimation, experimental, and recovery periods. Mussels were acclimated to four temperature x food treatments, where the low tide, aerial temperature was ramped to either 20 or 30 °C and food rations of 0.25 or 1.25 % algae·g mussel dry weight^-1^ of Shellfish Diet™ 1800 (Reed Mariculture, Campbell, CA, USA) were provided daily. Food was pulsed into the tanks during high tide cycles at 1 h increments. The temperatures within the tanks were controlled by the internal body temperature of the mussel, mimicked using ‘Robomussels’ (Helmuth et al., 2016). During the daytime, low tide, the air temperature was ramped to the appropriate treatment (e.g., 20 or 30 °C) at a rate of 0.1 °C per minute. Nighttime temperatures were held at 18 °C, while water temperature during high tide was kept at 16 °C. Mussels were acclimated for 3 weeks and a ‘baseline’ sample (e.g., gill tissue) was collected for comparison at 1 h into the afternoon high tide (e.g., 13:00). The following day, mussels in all treatments were exposed to an acute aerial heat shock of 33 °C during the daytime low tide and returned to acclimation conditions. Additional sampling occurred at 1 h into the recovery period and 24 h (a 25-h recovery) later, to control for circadian and circatidal variation in the proteome (Elowe & Tomanek, 2021). Five mussels were collected at each time point and treatment group; gill tissue was flash frozen and stored at −80 °C. More details can be found in Fabela et al. (in review).

**Figure 1.**
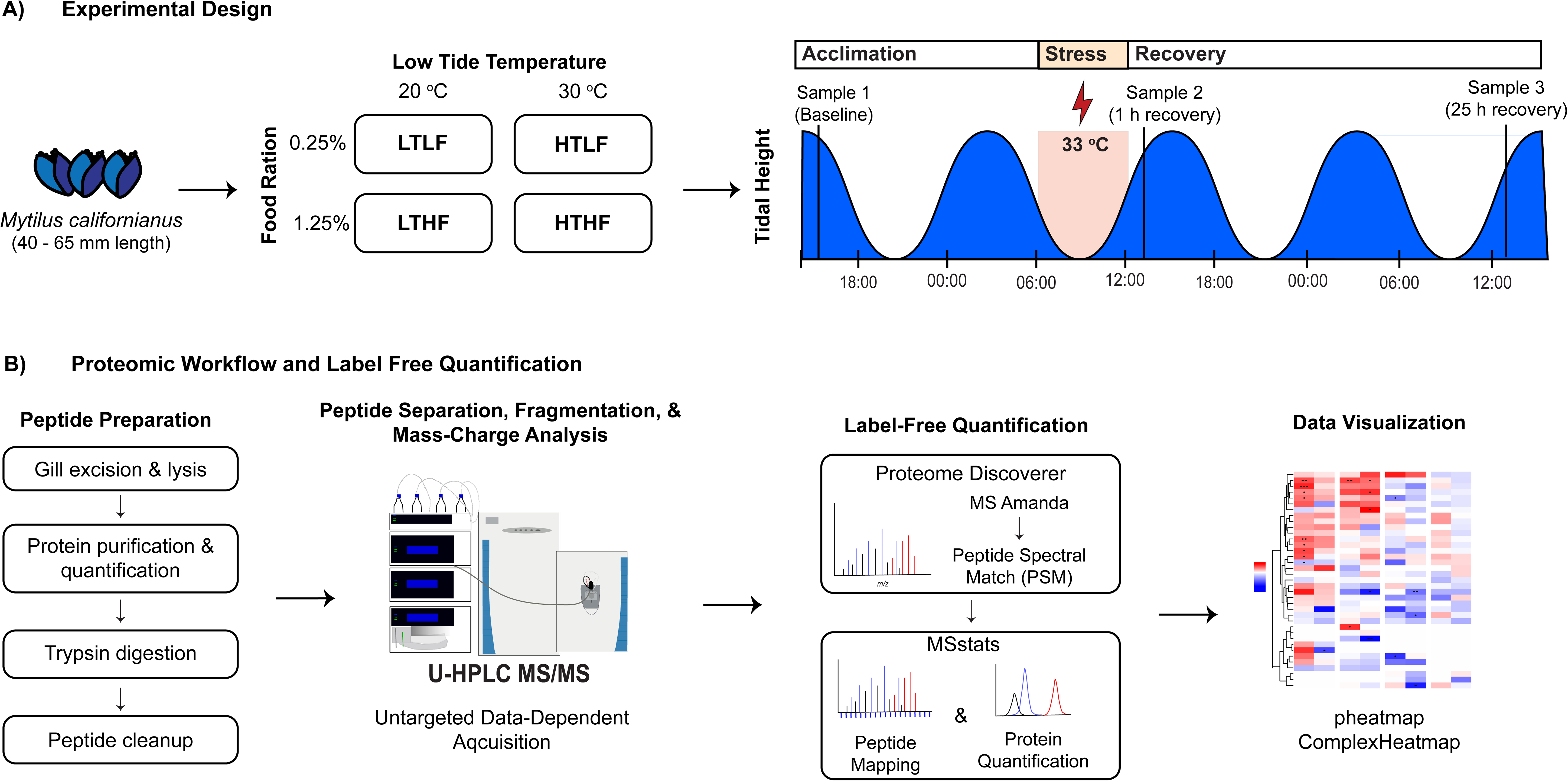
Experimental design and gill sample collections for *Mytilus californianus* heat shock exposure and recovery. The graphic shows A) the experimental design and B) proteomic workflow and label free quantification. Panel A includes the acclimation conditions of low or high temperature (20 or 30 °C) and low or high food ration (0.25 or 1.25% dry weight), corresponding to low temperature-low food (LTLF), low temperature-high food (LTHF), high temperature-low food (HTLF), and high temperature-high food (HTHF) groups, as well as, the timing of changes in water level, sampling events, and application of heat shock (33 °C). Panel B) overviews the proteomic workflow to analyze gill tissues using LC-MS/MS, including sample processing (peptide synthesis) and label free quantification to determine relative protein abundance of the recovery groups (1 h and 25 h past heat shock) compared to the baseline in MSstats (see Methods for details).

### Protein preparation and UHPLC-MS/MS analysis

Cell lysis was achieved via sonication of gill tissue (∼200 mg) in a denaturing buffer of 8 M urea (Thermo Scientific, Waltham, MA, USA), 2 M thiourea (Thermo Scientific, Waltham, MA, USA), and 1% deoxycholate (Spectrum Chemicals, New Brunswick, NJ, USA)(Feist & Hummon, 2015). Proteins were pelleted by centrifugation and cleaned overnight using 4x volumes of 10% trichloroacetic acid (Fisher Chemical, Pittsburg, PA, USA) in acetone (Fisher Chemical, Pittsburg, PA, USA). Proteins were again pelleted, washed in ice-cold acetone, and resuspended in 8 M urea with 150 mM Tris-HCl (Fisher Chemical, Pittsburg, PA, USA). We used Pierce™ BCA Protein Assay Kit (Thermo Scientific, Waltham, MA, USA) to determine total protein concentration prior to reduction and alkylation of proteins using dithiothreitol (4mM; Thermo Scientific, Waltham, MA, USA) and iodoacetamide (18 mM; BioRad, Hercules, CA, USA), respectively. Proteins were diluted to < 1M urea using ammonium bicarbonate (Thermo Scientific, Waltham, MA, USA) before digestion using Pierce™ trypsin protease (Thermo Scientific, Waltham, MA, USA). Before analysis using ultra-high-performance LC coupled with tandem MS (UHPLC-MS/MS), peptides were desalted using Pierce™ C_18_ spin columns (Thermo Scientific, Waltham, MA, USA). Separation and detection of peptides (run in triplicate) was conducted using an Ultimate 3000 and Q Exactive Plus Hybrid Quadrupole-Orbitrap MS (Thermo Scientific, Waltham, MA, USA). As described in Fabela et al. (in review), peptides were run through a 5 mm x 300 μm C_18_ precolumn (Thermo Scientific, Waltham, MA, USA) and a 150 mm x 75μm (particle size 2 μm) Acclaim PepMap 100 C_18_ HPLC analytical column (Thermo Scientific, Waltham, MA, USA) with a linear gradient of 1% buffer B (80% acetonitrile with 0.08% formic acid (FA); Fisher Chemical, Pittsburg, PA, USA) in buffer A (0.1% FA), to 35% buffer B over 90 min at a flow rate of 300 nl·min^-1^. The Nanospray Flex™ Ion Source had a spray voltage of 2.4 kV, a capillary temperature of 275°C, and S-lens RF level of 50. Full scans were taken between 375 −1575 *m/z* in positive mode at 70,000 resolution. The automatic gain control (AGC) target was 3e^6^, with a maximum injection time of 100 ms. MS/MS scans were taken with 17,500 resolution, an AGC target of 1e^5^, and a maximum injection time of 50 ms. The top 10 ions were fragmented at a normalized collisional energy of 27%. Dynamic exclusion was used with a duration of 20 s, the intensity threshold was set to 1.6 e^5^, with an isolation window of 2 m/z and lock masses turned on. We excluded analysis of peptides with charges of 1 and ≥7. All parameters were set using Thermo’s operational software (Tune v.2.12 build 3134 and XCalibur v. 4.4.16.14).

### Label Free Quantification

Initial processing of raw MS data was completed using Thermo Proteome Discoverer v.2.3 (PD), using processing and consensus workflows developed by (Griss et al., 2019). For peptide identification, we used MS Amanda with the *Mytilus galloprovincialis* (Lamark) (NCBI taxonomy ID: 29158) as a reference protein database and compared to a concatenated target decoy protein dataset with an FDR of p < 0.05 (Gerdol et al., 2020). Although a *M. californianus* genome database has since been published (Paggeot et al., 2022), its BUSCO completeness is below the former (∼86% versus 97.5% coverage of complete single-copy genes). Predicted proteins were searched allowing for trypsin digestion with 3 missed cleavages and tolerances of 7 ppm and 0.03 Da. Precursor and fragment ion tolerances were 5 ppm and 0.02 Da.

The processed peptide dataset was subsequently analyzed using MSstats R package v.4.6.5 (Choi et al., 2017). Peptides were matched to proteins using an FDR of 2% for unique peptides, peptides matched to one protein, and proteins with three or more matched peptides. Relative protein abundances were obtained by summing the max peak area of each matched peptide, applying a log_2_ transformation to the sums, and performing a median normalization across MS runs. Run-level summarization was performed using Tukey’s median polish. For model estimation, a linear mixed-effects model, with both biological and technical replicates included, was run, followed by whole plot, including the overall model median, the condition and subject by condition variation, the run variation, and a subplot with run variation and error term for each feature (i.e., peptide area) by run combination. Changes in protein abundance were determined by comparing Sample 2 (1 h following heat shock) and Sample 3 (25 h following heat shock to Sample 1 (the baseline), within each acclimation group (α = 0.05). The p-values were adjusted for multiple comparisons, and model assumptions were visually verified in MSstats; quality control was conducted using ProBatch (Cuklina et al., 2021). Visualization of the data via heatmap was performed using “pheatmap” (Kolde, 2025) and “ComplexHeatmap” (Gu et al., 2016).

## RESULTS

We identified over 1,000 gill proteins across our acclimation and treatment groups. Changes in protein abundance following heat shock are dependent on food-by-temperature acclimation (Table 1). Here, we present our synthesis of changes in abundance for proteins involved in mRNA-processing, protein synthesis, molecular chaperones, protein degradation, actin- and microtubule-associated cytoskeleton, the extracellular matrix, and cellular signaling. The role of food in modulating the CSR with respect to core carbohydrate and one-carbon metabolism, oxidative stress, and DNA repair has been discussed in a companion manuscript (Fabela et al., in review). Consistent with our previous findings, mussels acclimated to low food ration and lacking thermal conditioning (e.g., LTLF) mounted a larger cellular response, suggesting significant need for maintaining protein and cellular structure. We therefore focus much of the results and discussion on the response in LTLF before describing changes across treatments.

**Table 1.**
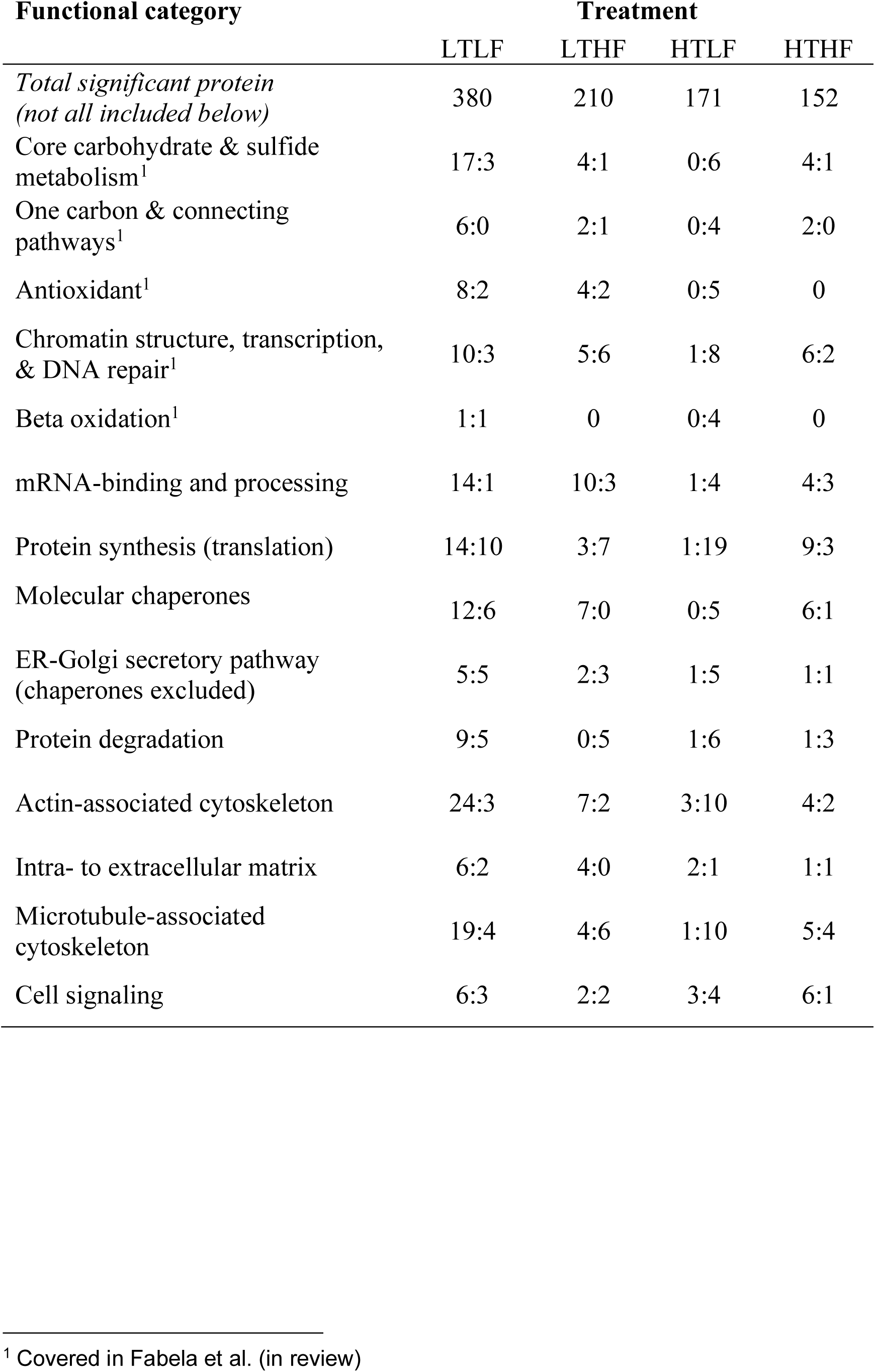
Changes (increases:decreases) in protein abundance following acute heat stress by food-by-temperature acclimation treatment included in heat maps.

### mRNA-binding and processing proteins

We identified several small nuclear ribonucleoparticle (snRNP) proteins, splicing factors, heterogeneous nuclear ribonucleoproteins (hnRNPs) and RNA-binding proteins (RBPs) that associate with nascent transcripts (Lodish et al., 2021; Marasco & Kornblihtt, 2023), some of which are essential for alternative splicing during heat stress (Haltenhof et al., 2024). While selective changes in mRNA processing proteins occurred in all but one acclimation group (high-temperature-high food – HTHF), we observed additional increases in low-temperature-low-food (LTLF), and a similar but delayed response in low-temperature-high-food (LTHF) acclimated mussels (Fig. 2), suggesting food-dependent temporal heterogeneity within LT-acclimated mussels. In either group, several arginine/serine-rich splicing factors (SRSF1/9, 4/5/6, 7, and 16) increased, which direct spliceosome assembly to generate mRNA variants tailored to stress conditions. They also promote the recruitment of snRNPs (e.g., U5 snRNP) to control alternative splicing, thereby mediating nuclear export and translation efficiency and linking splicing with gene expression regulation (Cho et al., 2011; Shen & Mattox, 2012). SRSF7 is specifically involved in splicing regulation under stress conditions, including oxidative and heat stress causing DNA damage (Haltenhof et al., 2024), and two isoforms increased in LT-acclimated mussels. The hnRNPs are involved in different stages of mRNA processing, including splicing, mRNA stability, and translational and transcriptional regulation (Geuens et al., 2016; Shi et al., 2017). LTLF mussels also increased three hnRNPs (A1/A3, F/H, R, and U-like), the translational regulator RNA-binding protein Musashi (Banik et al., 2024), a “reader” of methyladenosine-modified RNA YTH-domain family protein (Shi et al., 2021), and Y-box-binding protein 1, a transcription factor associated with stress granules and translation regulation (Maurya et al., 2017), emphasizing the broad spectrum of possible RNA processing modifications during acute heat stress.

**Figure 2.**
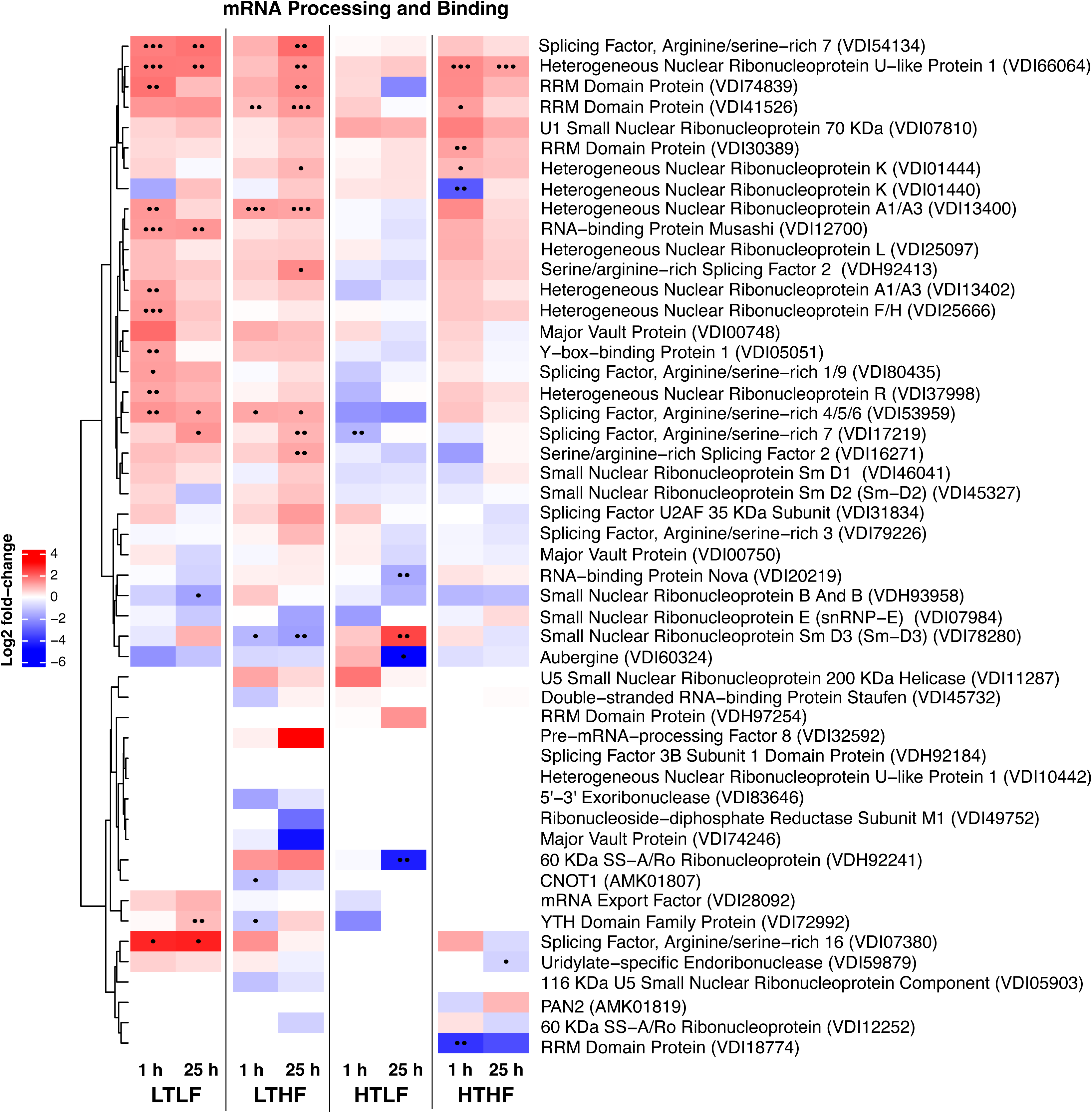
Heat map of mRNA-binding and processing proteins. The heat map shows log_2_ fold change in protein abundance from 1 h or 25 h recovery from heat shock relative to the baseline in each of the acclimation groups: low temperature-low food (LTLF), low temperature-high food (LTHF), high temperature-low food (HTLF), and high temperature-high food (HTHF). Increases in abundance are indicated by warm colors (reds), while decreases are cool (blues). Significance levels are shown by dots, where ••• signifies p < 0.001, •• is p < 0.01, and • is p < 0.05. Proteins were identified using GenBank accession numbers.

LTHF also increased several SRSF and hnRNPs, several of which overlapping with LTLF mussels. In contrast, HT-acclimated mussels showed fewer changes overall, especially in HTLF (Fig. 2). Across acclimation conditions, snRNP proteins were relatively stable following heat stress, but the nucleolar hnRNP U-like 1 increased in all but HTLF, possibly affecting ribosome biogenesis (Cichocka et al., 2022). Thus, LT-acclimated mussels increased a broad spectrum of RNA-binding proteins, although delayed in LTHF, while HT-acclimated mussels changed almost none (HTLF) or far fewer (HTHF).

### Protein Synthesis

Protein synthesis is estimated to require a substantial part of the cell’s energy consumption (Rolfe & Brown, 1997), and is likely finely tuned to food intake. Several proteins related to protein synthesis, including the ribosome, translation, mRNA structural stabilization, and aminoacyl-tRNA ligation changed following heat shock (Fig. 3), particularly in the low food and temperature (LTLF) mussels (Table 1). Ribosomal proteins (RPs) accounted for most of the observed changes, as did two translation initiation factor (eIF3G, 3L) subunits, which control translation rates of some mRNA by scanning regulatory codes present in the 5’ UTR (Choudhuri et al., 2013). Two elongation factors (eEF) increased eEF1α in both LT-acclimated mussels and eEF2 only in LTLF; eEF2 serves a regulatory role, where its phosphorylation inhibits translation (Holcik & Sonenberg, 2005). Lastly, of the five aminoacyl-tRNA ligases, two decreased in LTLF while two others showed opposite responses in LTHF mussels. Although reduced during stress in LTLF, asparaginyl-tRNA ligase presents a node connecting amino acid availability, tRNA homeostasis, and stress-adaptive translational control, and its tRNA charging capacity remains essential to stress granule and P-body assembly for RNA sequestration (Emara et al., 2010). Thus, LTLF mussels modified translation by mostly increasing selective RPs, translation initiation and elongation factors, and amino acid-tRNA ligases. Some protein decreases indicate translational dysregulation.

**Figure 3.**
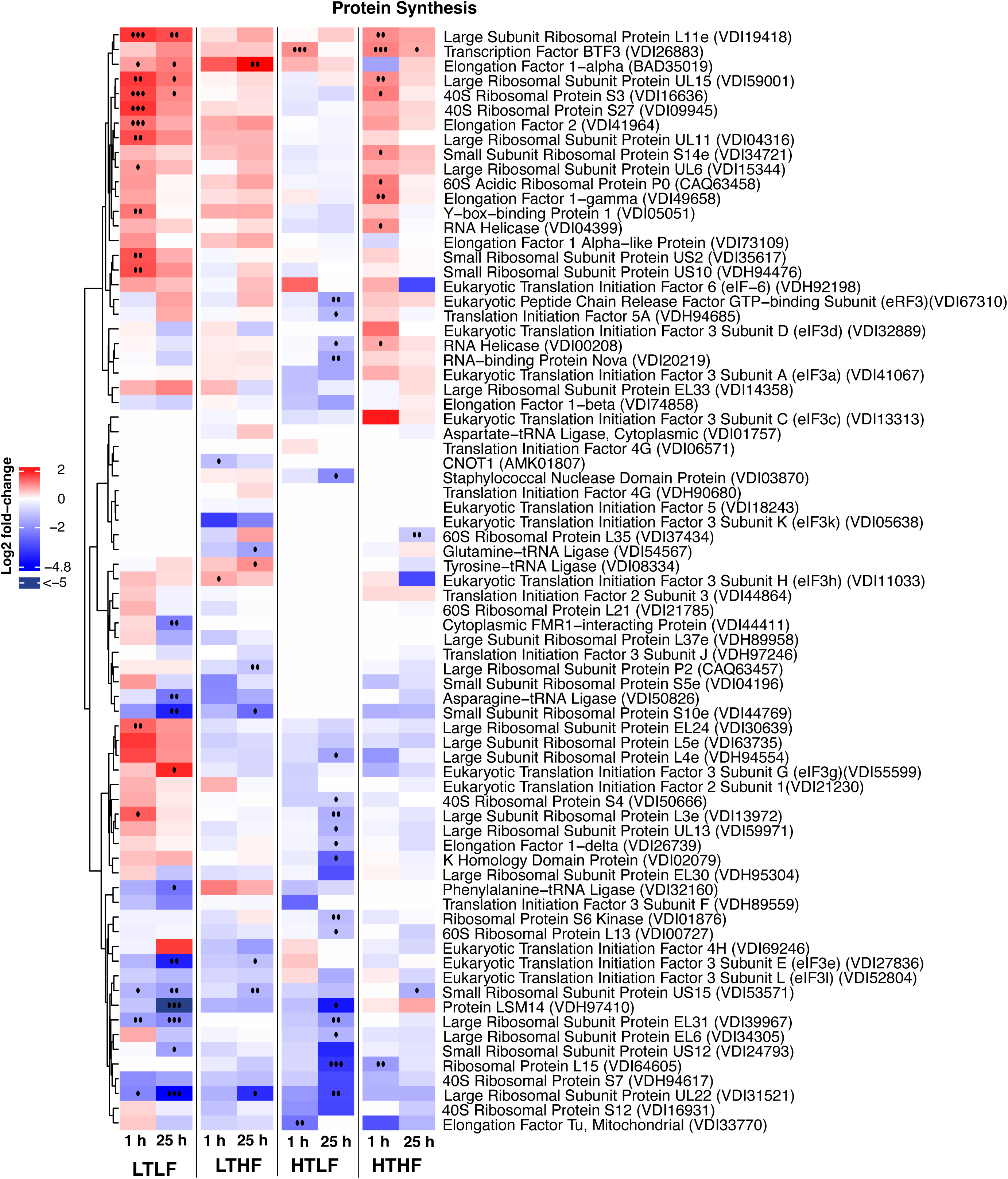
Heat map of ribosome and translation proteins. The heat map shows log_2_ fold change in protein abundance from 1 h or 25 h recovery relative to the baseline in each of the acclimation groups; For additional details see Fig. 2.

In comparison, low-temperature-high-food (LTHF) mussels changed few translational proteins. In contrast, high-temperature-low-food (HTLF) mussels decreased nineteen translational proteins, only three of which also decreased in LTLF. And although HTHF increased nine RPs, only three overlapped with those increasing in LTLF. The overall translational response was therefore fine-tuned to each acclimation condition. Unique to HTHF mussels, two RNA helicases, which stabilize RNA secondary structure during heat stress to maintain translation and are involved in the assembly of stress granules to sequester non-translating mRNA during stress, increased with heat (Owttrim, 2006). Overall, ribosomal modifications and translation initiation and elongation are important for shifting the acclimation-specific baseline translational machinery in response to acute heat stress.

### Molecular Chaperones

#### Cytosolic chaperones

Heat shock inhibits translation of housekeeping genes, requiring a selective and quick increase of stress proteins to maintain proteostasis (Morimoto, 1998). Of the 37 proteins we categorized as molecular chaperones, nearly half changed (12 increased; 6 decreased) in low-temperature-low-food (LTLF) mussels (Table 1; Fig. 4). These data suggest that LTLF mussels experienced a greater proteostasis challenge than other groups. Heat shock proteins (Hsps) are the main molecular chaperones, responsible for protein chaperoning and folding following heat and other environmental stressors (Somero et al., 2017). We observed increases in the abundance of a cytosolic and endoplasmic reticulum Hsp70s (HspA5)—the main chaperone family capable of both housekeeping and stress-related activities—in LTLF mussels. Another decreased in LTLF, but this isoform is homologous to Hsp12A, which is involved in regulating aerobic glycolysis following oxygen reperfusion in cardiac muscle and not heat-related responses (Li et al., 2024), a process analogous to the onset of high tide in the present study. While these chaperones possess a broad target spectrum, other chaperones have more specific client proteins. For example, heat shock protein 90 plays a key role in several multi-chaperone complexes thereby affecting signaling pathways (Li et al., 2012). Hsp90 isoforms can be constitutively expressed at high levels (e.g., 1-2% of protein pool) or increase in response to cellular stress to ensure survival under acute stress (Bhattacharya et al., 2022). Hsp90 increased in LTLF and high-temperature-high-food (HTHF) mussels.

**Figure 4.**
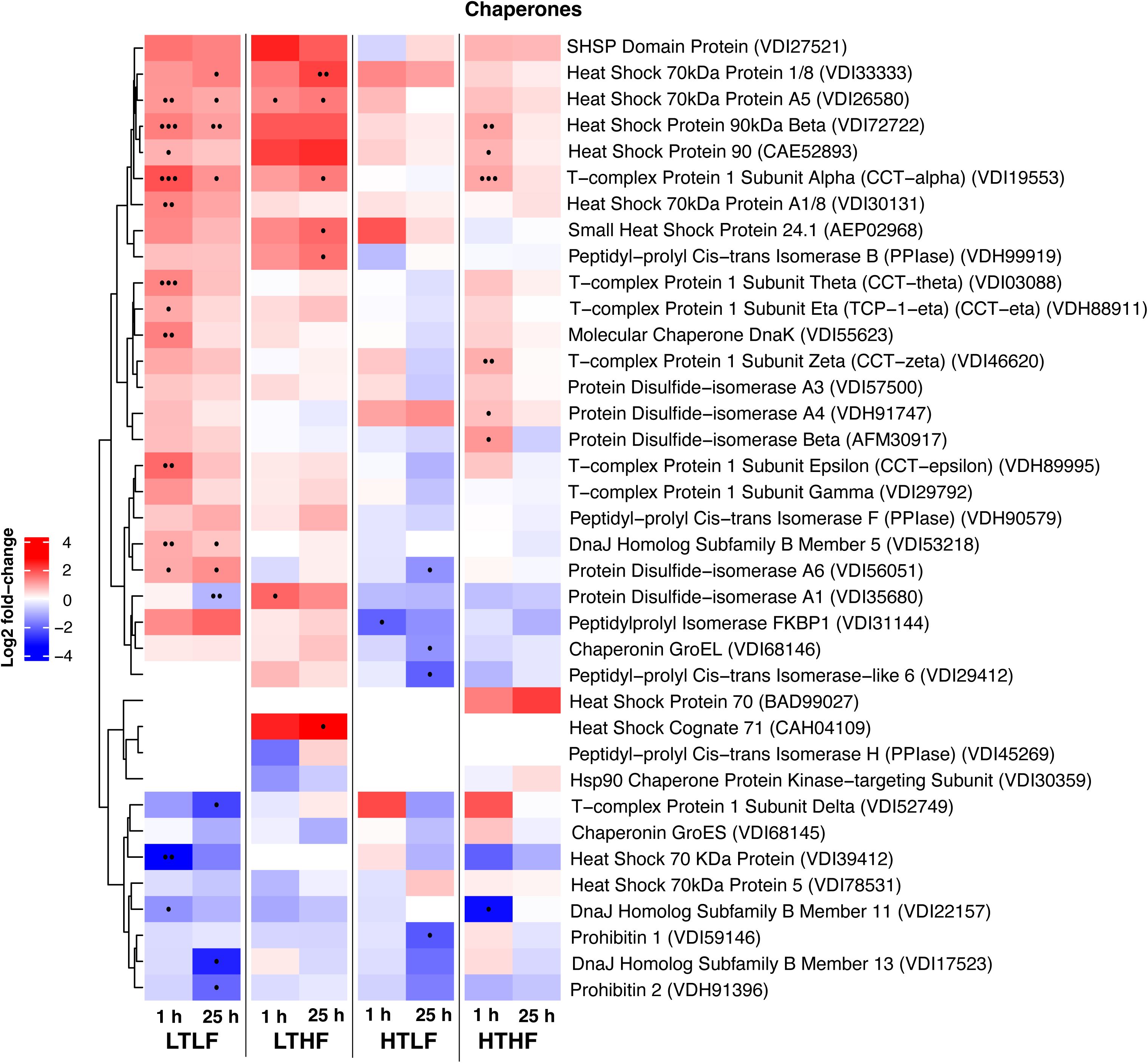
Heat map of molecular chaperones. The heat map shows log_2_ fold change in protein abundance from 1 h or 25 h recovery relative to the baseline in each of the acclimation groups; For additional details see Fig. 2.

DnaJ (Hsp40) co-chaperones enhance Hsp70’s ATP-hydrolyzing activity and target specificity (Hartl et al., 2011; Qiu et al., 2006). Of three DnaJ isoforms we identified, the cytosolic DnaJB5 increased in LTLF and stayed elevated at 25 h, while the ER-localized DnaJB11 decreased, possibly to activate the unfolded protein response (UPR) by freeing up an ER Hsp70 homolog (see below). DnaJB13, which is localized to the central apparatus of cilia, also decreased, possibly affecting cilia stability. We also observed an increase in the mitochondrial chaperones DnaK and a decrease in prohibitin 2 in LTLF mussels. While small Hsp (sHsp) have been implicated in interspecific differences in thermal tolerance and protecting cells from heat and oxidative stress through their ability to hold and stabilize unfolded proteins (Tomanek & Zuzow, 2010), we surprisingly did not observe significant increases in LTLF, but did in low-temperature-high-food (LTHF) mussels. See Vasquez et al. (in review) for how mussels from a separate experiment, but under conditions similar to our baseline acclimation groups, differed in sHsp abundance.

Finally, we observed increases in the chaperone-containing tailless complex polypeptide 1 (CCT-α, −ε, −η and −θ), also known as T-ring complex (TRiC), in LTLF mussels. CCT assists in folding predominantly cytoskeletal proteins, such as actin and tubulin (Vallin & Grantham, 2019), indicating that LTLF mussels require greater folding activity of cytoskeletal proteins as we provide evidence for below. Other acclimation conditions varied other CCT-subunits, possibly due to the association of their monomeric forms with microtubules (CCT-α,-ζ and −η), actin bundles (CCT-ε), and actin-severing and -capping gelsolin performing moonlighting functions (Brackley & Grantham, 2010; Roobol et al., 1999; Svanstrom & Grantham, 2016).

#### Endoplasmic Reticulum (ER) chaperones and secretory pathway proteins

As native polypeptides emerge from the ribosome, they are translocated into the ER lumen and pulled in by glucose-regulated protein (Grp78, or BiP), which initiates folding by additional chaperones, and can serves as an ER stress sensor to activate the ER unfolded protein response (UPR; Fig. 4; Braakman & Hebert, 2013). These proteins increased predominantly in LT-acclimated mussels, indicating a need to activate the UPR. This is further supported by decreases in the Hsp40 co-chaperone DnaJB11, which increases HspA5’s (the Grp78 homolog) ATP-hydrolyzing activity while independently binding protein targets and transporting them across the ER (Hartl et al., 2011; Qiu et al., 2006). Thus, HspA5’s decrease may facilitate the UPR (Larburu et al., 2020). We also observed an increase in the ER-localized protein disulfide isomerase (PDI) in LTLF mussels. PDIA6 attenuates ER stress sensor IRE1a signaling (Eletto et al., 2014) and its increase dampens the magnitude and duration of UPR signaling, suggesting that LTLF mussels tried to maintain folding glycoproteins by ER chaperones while simultaneously limiting the UPR. Increases in PDIA6 were not observed in low-temperature-high-food (LTHF) mussels, likely because increased food rations can induce higher constitutive levels of PDIA6 and the UPR (Deldicque et al., 2010). Other PDI isoforms involved in disulfide bond formation and protein folding (Bulleid, 2013), varied in other acclimation groups (Fig. 4).

Evidence for an acclimation-specific ER proteostasis challenge also comes from peptidyl prolyl *cis-trans* isomerases (PPIases), which assist in proline isomerization, a rate limiting step in ER protein folding (Braakman & Hebert, 2013). PPIases interact with Grp78 and enhance its chaperone functions; of five PPIase isoforms, one increased in LTHF at 25 h, while two decreased in high-temperature-low-food (HTLF) mussels. After folding, the next maturation step involves a multimeric oligosaccharyl transferase complex, which commits the protein to glycosylation in the *cis*-Golgi and increased in LTLF, which are presented in a separate figure from the ER chaperones (Fig. S1). Further processing involves trimming oligosaccharides by glucosidase II, quality control chaperoning by calnexin, and the stepwise degradation of glycoconjugates in lysosomes (Lodish et al., 2021). Many vesicle-associated proteins were identified, including coat proteins and small G-proteins involved in *cis*-Golgi to ER retrograde transport (COPI-associated Sec 22/23/24, P24, and Sar1), *trans*-Golgi to endosome (clathrin-AP-1 complex and Arf) transport, vesicle docking to compartment specific membranes (Rab isoforms), and vesicle pinching (dynamin). Several of these tended to be lower (mostly non-significantly) across treatments, especially in LTHF and HTLF (Fig. S1). In addition, increased atlastin, an ER structural protein functioning in ER membrane fusion, suggests damage to overall ER structural integrity and may indicate ER-autophagy in HTHF (Fig. 4; Liang et al., 2018). This conjecture is supported by a decrease in vesicle-associated membrane protein-associated protein A (VAPA) which helps maintain ER membrane tethering to other organelles (James & Kehlenbach, 2021) and has also been linked to autophagy (Zhao et al., 2018).

In HTLF mussels we observed a different trend, with decreases in protein folding proteins, such as PDIA6 and two PPIs (FKBP1/6), indicating that protein maturation in the ER was suppressed following heat shock (Fig. 4). Additionally, glucosidase IIα, along with vacuolar protein-sorting-associated protein 4 and the small G-protein ADP-ribosylation factor 8 (Arf 8) which regulates vesicular trafficking, with the latter decreasing in all but HTHF (Fig. S1). These support a lack of a response or even a downregulation of ER protein maturation across acclimation conditions (Table 1).

The oligosaccharides attached to secretory proteins originate with the hexosamine pathway, a branch of glycolysis, including glucosamine-fructose-aminotransferase (Gfat), phospho-acetylglucosamine mutase (Pagm), and UDP-acetylglucosamine diphosphorylase (Uap; Chandel, 2015), and are essential building blocks for glycosylation. We observed an increase in Gfat in LTLF, the rate-limiting step, and in Pagm at 25 h in LTHF, but a decrease in Uap after 25 h in LTLF, suggesting an acclimation specific response (Fig. S1). We also identified proteins involved in mannose biosynthesis, consistent with the finding that mucopolysaccharides of mucus samples are composed mostly of mannose residues (Pales Espinosa et al., 2016), as well as enzymes within fucose biosynthesis, and with hydrolysis and transferase activities.

### Protein Degradation Pathways

Following iterative attempts to stabilize misfolded proteins and ordering of partially misfolded proteins into protein aggregates, chaperones also assist in delivery of irreversibly denatured proteins to the proteolytic machinery, to prevent deleterious effects and recycle amino acids. We detected proteins of several stages of the ubiquitin-proteasomal-system and small amino peptidases (Fig. 5; Ciechanover, 2005). Overall, few degradation-related proteins changed significantly, and we almost exclusively observed increases in protein abundance in the low-temperature-low-food (LTLF) mussels (Table 1).

**Figure 5.**
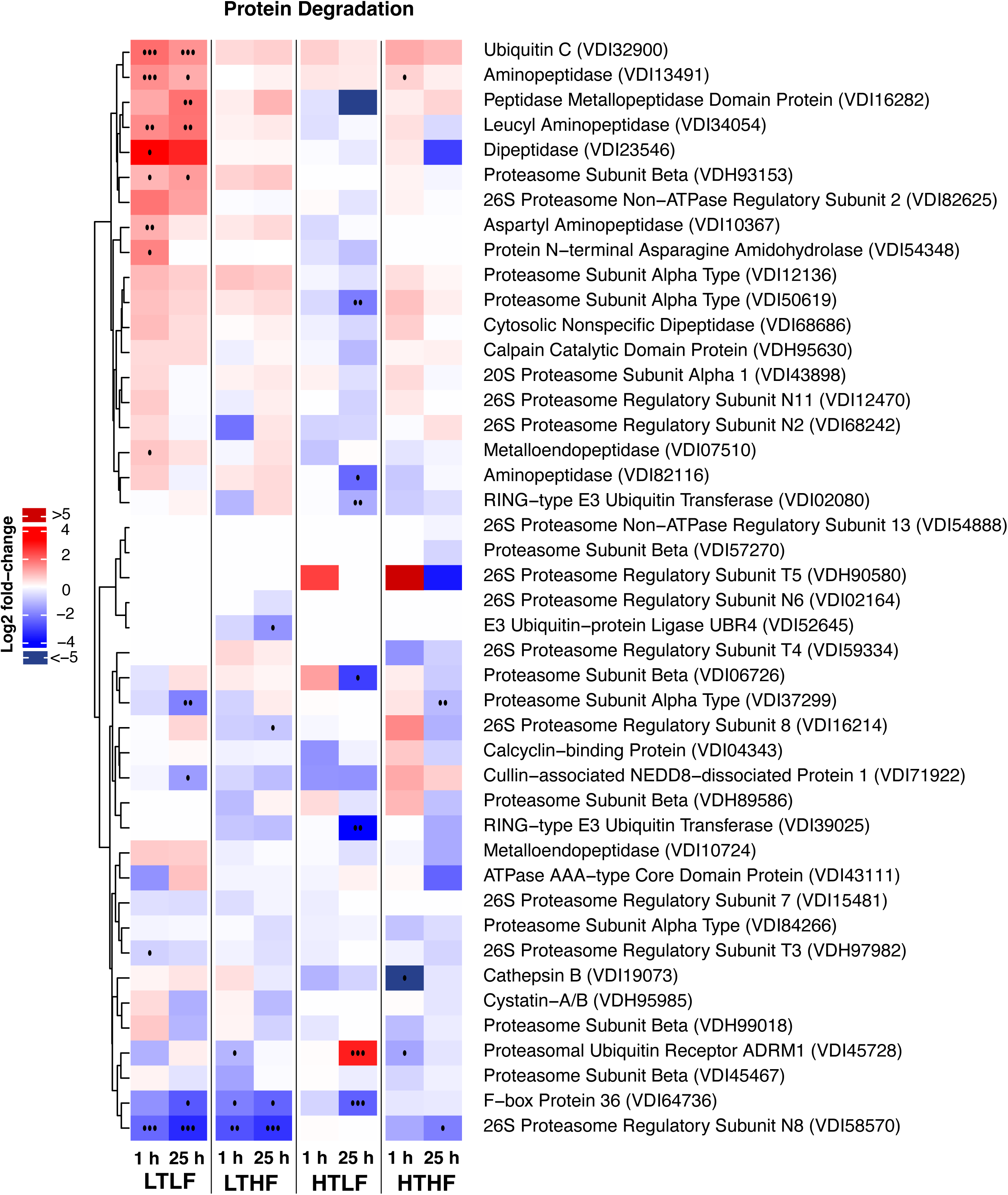
Heat map of protein degradation proteins. The heat map shows log_2_ fold change in protein abundance from 1 h or 25 h recovery relative to the baseline in each of the acclimation groups; For additional details see Fig. 2.

At the heart of protein degradation is the 26S proteasome, consisting of the 20S core particle, which is responsible for the proteolytic activity, and the 19S regulatory particle, which recognizes ubiquitin-labeled proteins, unfolds them, and guides them into the proteolytic chamber—a process requiring ATP (Finley, 2009). The increase of proteasome β subunit and decreases of 26S proteasome regulatory subunit T3, an ATPase that facilitates the opening of the 20S core particle to allow the unfolded substrate to enter, is potentially evidence for modifications of UPS activity. LTLF also increased ubiquitin C, which is required for ubiquitination. Furthermore, the decrease of several additional components suggests patterns of ubiquitination immediately shift following an acute heat stress. Two substrate-recognition components of E3 ubiquitin ligase complexes (Cullin-associated NEDD8-dissociated 1 and F-box 36), a regulatory subunit that recognizes ubiquitinated proteins (subunit N8), and another that facilitates the opening of 20S core particle (α subunit), all decreased. The proteasomal ubiquitin receptor ADRM (also known as Rpn13), a non-ATPase subunit of the 19S regulator base responsible for recognizing di-ubiquitin and recruiting of protein substrates, didn’t change in LTLF, but increased at 24 h in HTLF, and decreased in HF-acclimated mussels, indicating that it’s response to heat stress is unique to each acclimation condition. Related to degradation, LTLF mussels increased N-terminal asparagine aminohydrolase, a protein responsible for catalyzing the deamidation of N-terminal asparagine to aspartate, which controls protein lifespan and turnover (Varshavsky, 2019), providing further evidence for extensive shifts in processing of irreversibly damaged proteins and protein turnover in this group.

Following proteasomal degradation, the resulting peptides are further degraded by aminopeptidases, a family of proteins cleaving amino acids from the peptides’ N-terminus. Leucyl aminopeptidase, which increased in LTLF, hydrolyzes these peptides and glutathione into small fragments, thus facilitating glutathione turnover (Josch et al., 2003). As previously reported (Fabela et al., in review), LTLF mussels appear to experience high levels of oxidative stress, employing the tripeptide glutathione and other antioxidant systems to restore redox balance. Thus, increases in protein degradation in LTLF may serve complimentary roles: degradation of irreversibly damaged proteins and facilitating amino acid turnover for new proteins, including glutathione, one of the most abundant cellular metabolites and a non-enzymatic antioxidant. Glutathione may also be needed for the synthesis of lipid messengers, i.e. prostaglandins (see below).

### Cytoskeletal Proteins

#### Actin Filament Growth and Stabilization

Actin filaments provide structural integrity, function in intracellular transport, and cell-cell or cell-ECM adhesion (Pollard, 2016). Low-temperature-low-food (LTLF) mussels increased actin filament capping (actin-capping protein/CapZ, tropomyosin, but not enabled, tropomodulin), nucleation (two Arp2/3, but not formin/plastin), severing (gelsolin, advillin), and disassembling proteins (actin-interacting protein 1/Aip1), but not proteins involved in actin treadmilling (e.g., profilin, cofilin, costars, thymosin), suggesting that cells stabilized existing filaments, nucleated new branched filaments, and disassembled others following heat shock, possibly to augment filament growth elsewhere (Fig. 6).

**Figure 6.**
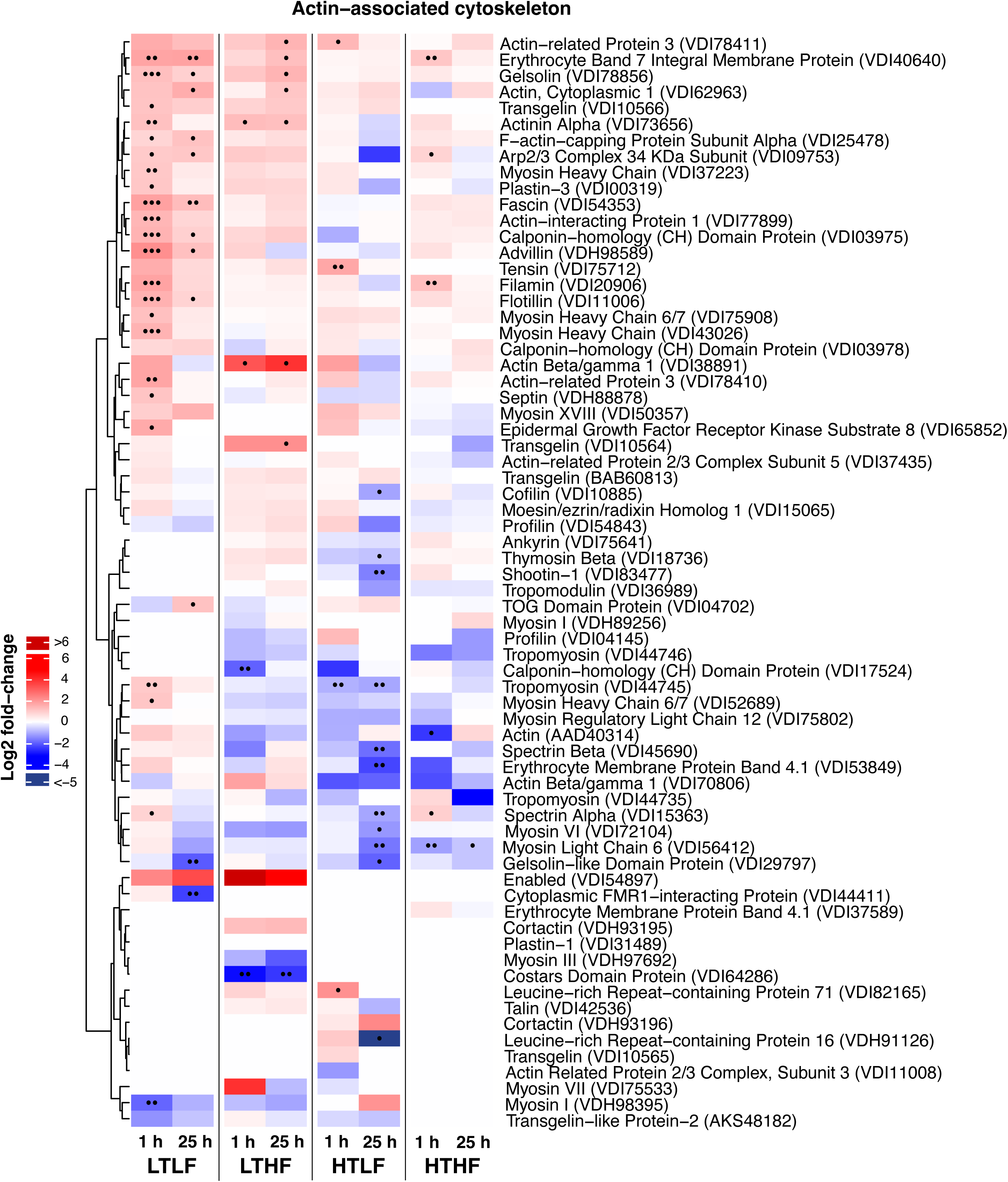
Heat map of actin-associated cytoskeletal proteins. The heat map shows log_2_ fold change in protein abundance from 1 h or 25 h recovery relative to the baseline in each of the acclimation groups; For additional details see Fig. 2.

Actin binding proteins (ABPs) stabilize and connect filaments by cross-linking or bundling filamentous actin. LTLF mussels increased ABPs that contribute to stress fiber formation (α-actinin, filamin, fascin, transgelin, and calponin-homology (CH) domain-containing protein isoforms) and facilitate formation of a scaffold-like structure that strengthens the cell cortex by connecting to the cell membrane (spectrin α, flotilin), or join filaments to focal adhesions, attaching the cell to the ECM (plastin-3/formin; Pollard, 2016). ABPs attaching the filaments directly to the membrane did not change (e.g., erythrocyte membrane protein band 4.1, ankyrin, moezin/ezrin/radixin). However, we did observe an increase in erythrocyte band 7 isoform (or stomatin) and septin, which localize to membrane protrusions and maintain cell shape, as well as myosin isoforms (myosin heavy chain 6/7), all of which are likely involved in anchoring filaments to the membrane and vesicle transport (Sweeney & Holzbaur, 2018). Thus, LTLF mussels respond by stabilizing actin and creating stress fibers to potentially modulate cytoskeletal and membrane tension (Katsuta et al., 2024).

Stress fibers connect the ECM through several ABPs, (talin, vinculin, afadin actinin) via transmembrane integrins (α6, α8, α9, and β) that are modified by kindlin. These complexes connect to type IV collagens (α-type), laminins, the cross-linking perlecan and the lattice-forming gelsolin to form the basal lamina (Fig. S2; Bachir et al., 2017; Lodish et al., 2021), which increased in LT-acclimated mussels. LTLF also increased two collagen types (XVIIIα and C4-like) that are integral membrane proteins associated with hemidesmosomes, while decreasing fibronectin type-III and integrin β, both of which play a key role in sensing rigidity of the ECM (Lodish et al., 2021). In comparison to LTLF, other acclimation groups showed a limited response.

Low-temperature-high-food (LTHF) mussels elevated ABPs, e.g., α-actinin and transgelin, typical for stress fiber formation (Fig. 4). In HT-acclimated mussels, the observed increases of different Arp2/3 isoforms across groups suggests enhanced actin nucleation during heat shock, accompanied in HTLF by an early increase followed by a decrease in leucine-rich repeat-containing protein 71 and 16 (or capping protein Arp2/3 myosin I linker/CARMIL, but not cortactin), respectively, which modulates actin assembly at branched networks (Jung et al., 2001). Overall, the muted response of HT-acclimated mussels suggests that ABPs stabilizing the cytoskeleton coped better with heat stress compared to LTLF. Collectively, these findings show that the stabilization of actin filaments is graded according to the overall magnitude of the CSR.

#### Microtubules

The microtubular (MT) system provides ciliary structure and, with its associated motor proteins, is responsible for organelle and intraflagellar transport. In mussels, lateral cilia generate the water flow across the gills, while frontal cilia transport food particles to the mouth, consuming >90% of the respiratory demand of the gill (Clemmensen & Jøgensen, 1987). Thus, we assume that most of the eight-nine MT-associated proteins, including nineteen cilia- and flagella-associated proteins (CFAPs), are associated with cilia structure and function (Fig. 7). Tektin-1, −3, and −4 that make up a core bundle inside the A-tubule, and MT-associated protein 1 (MAP1), echinoderm MAP 1/2, and CFAP52 that stabilize MT doublets along the axoneme and can link them to actin filaments, increased in LTLF (Conkar & Firat-Karalar, 2021; Gu et al., 2023). Other proteins that increased contribute to MT dynamics (tubulin polymerization-promoting 3), stability of cilia against mechanical stress (rootletin), proper docking of the outer dynein arm (outer dynein arm-docking complex), coordination of ciliary beating (hydrocephalus-inducing protein/hydin, CFAP57/61, radial spoke head 1/9) or are core inner MT-proteins (CFAP45/77). As components of the inner dynein arms and nexin-dynein regulatory complex, the proteins support ciliary motility and facilitate cilia so they can beat in a coordinated way (Bustamante-Marin et al., 2020). While all other acclimation groups showed half the number of proteins changing, HTLF mussels were different in that they mainly decreased MT-associated proteins.

**Figure 7.**
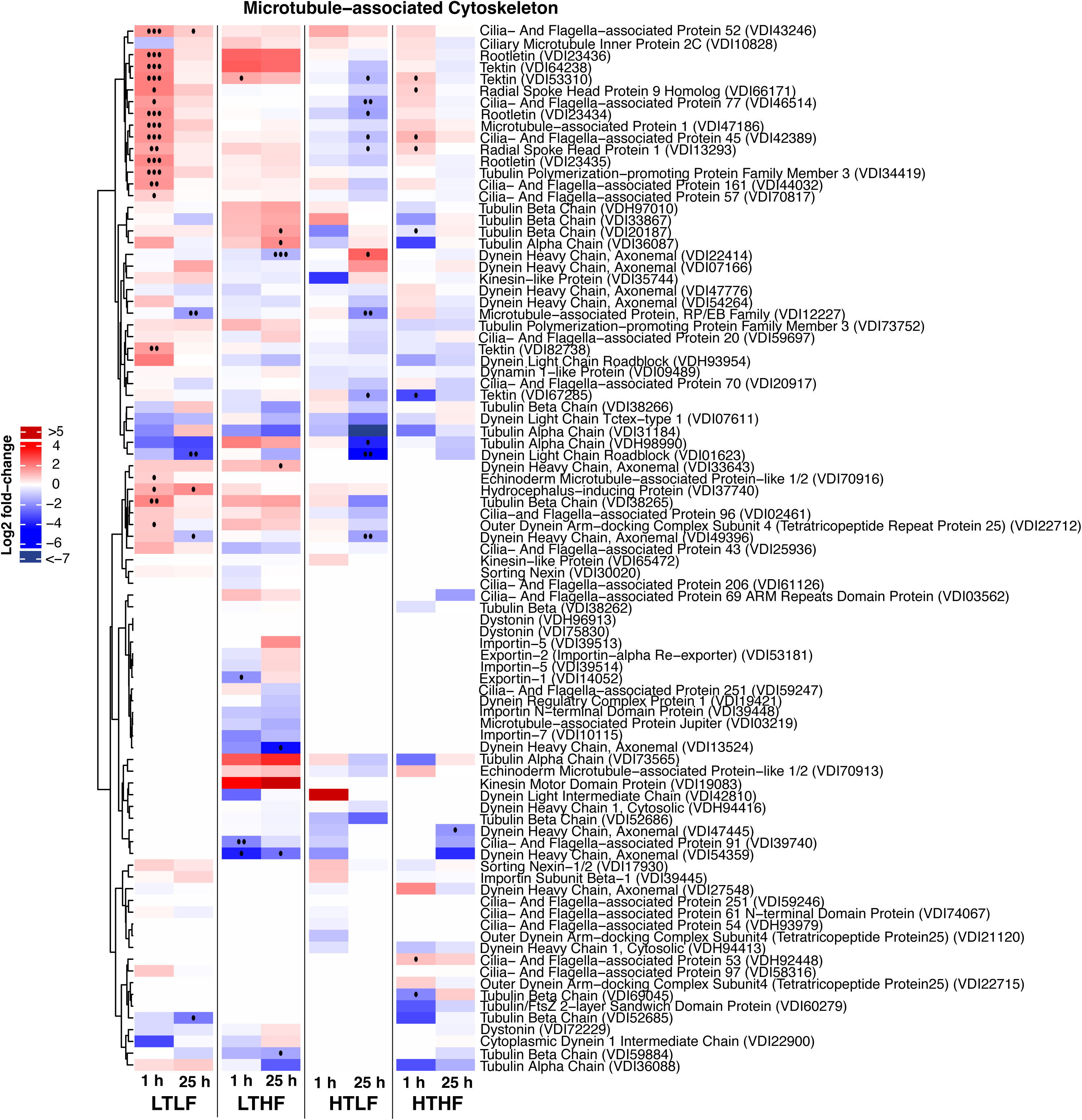
Heat map of microtubule-associated cytoskeletal proteins. The heat map shows log_2_ fold change in protein abundance from 1 h or 25 h recovery relative to the baseline in each of the acclimation groups; For additional details see Fig. 2.

### Cellular Signaling Proteins

Although signaling pathways mainly rely on post-translational modifications and changes in cellular location, abundance changes of signaling proteins also indicate heat-induced adjustments in these pathways (Fig. 8; Marks et al., 2017). Further, protein identification can identify signaling pathway typical for gill cells. For example, we identified several small G-protein families or their GTPase-activating proteins (Ras, Rho, Rab, Arf (Sar1), Ran) that represent multiple cellular processes, regulating cell survival and apoptosis, cytoskeletal dynamics, ER-Golgi secretory, nuclear and intraflagellar transport (Lodish et al., 2021; Marks et al., 2017). The Ras homolog Kras activates several downstream pathways, including the MAPK pathways, and can inhibit stress-induced factors of the ER UPR (Lv et al., 2023). Arf/Sar1 represents another Ras superfamily that plays a role in regulating the ER UPR and the transport of proteins from the ER to Golgi compartments under stress (Van der Verren & Zanetti, 2023). Arf/Sar1 members do this by initiating COPII-type vesicle formation to move unfolded proteins out of the ER for degradation. A Ras GTP-activating protein (IQGAP1), which primarily downregulates Ras, also contributes to the regulation of actin dynamics and the reorganization of the cytoskeleton by binding directly to F-actin and interacting with proteins involved in cytoskeletal reorganization (Brandt & Grosse, 2007). Three of these small G-proteins decreased, two in LTLF (Arf8 and an unassigned small GTPase) and one in HTHF (Arf-GTPase-activating 2/3).

**Figure 8.**
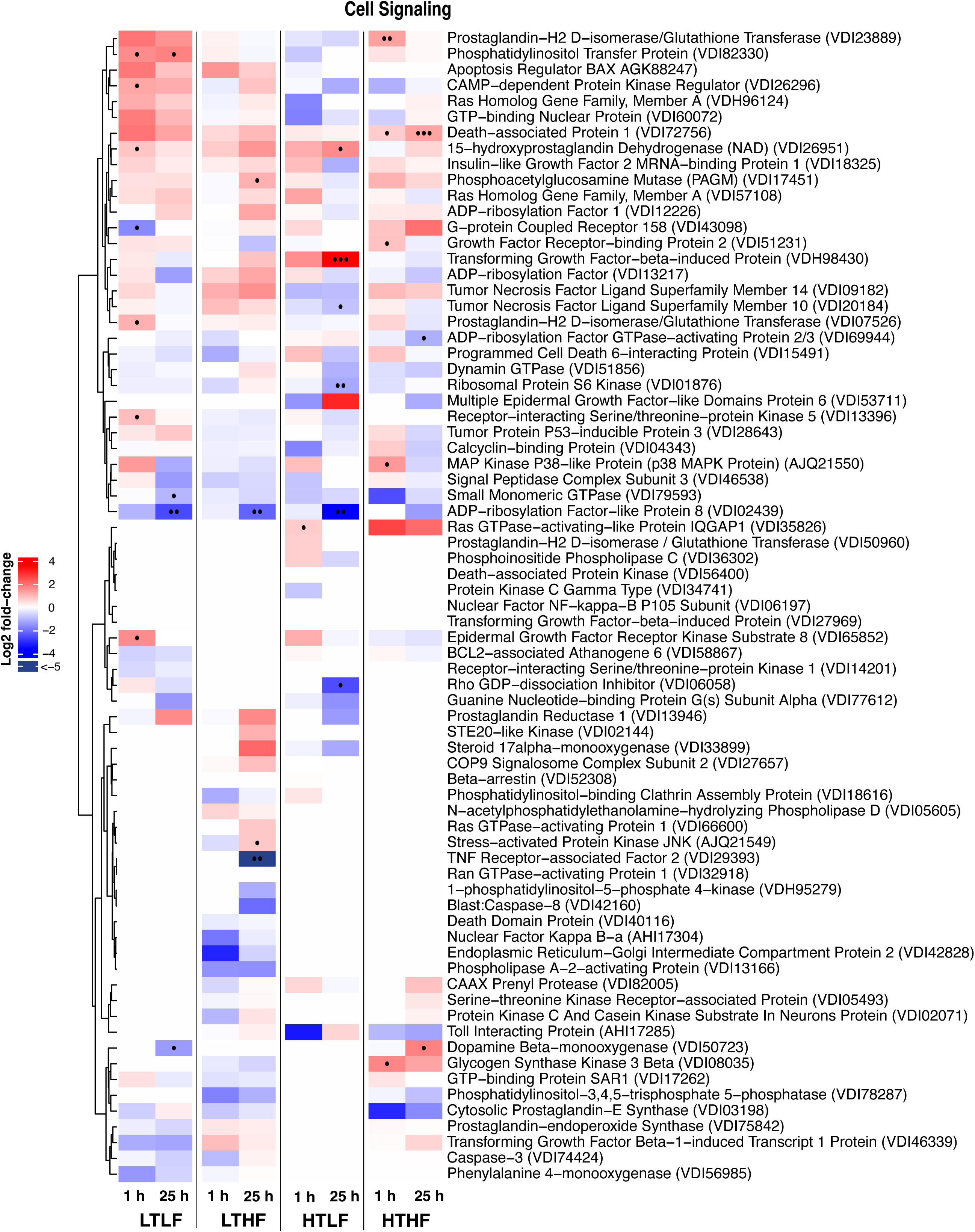
Heat map of cell signaling proteins. The heat map shows log_2_ fold change in protein abundance from 1 h or 25 h recovery relative to the baseline in each of the acclimation groups; For additional details see Fig. 2.

A select few signaling proteins increased in low-temperature-low-food (LTLF). An increase in receptor-interacting Ser/Thr-kinase 5 (RIPK5), which controls necrosis and serves as scaffolding protein for cell proliferating and apoptotic signaling proteins, is consistent with activation of cellular stress and DNA damage responses in LTLF (Fabela et al., in review; Zha et al., 2004). Heat shock also increased epidermal growth factor receptor kinase substrate 8 (EPS8), a signal adapter protein in the epidermal growth factor receptor (EGFR) pathway. By binding to actin, EPS8 facilitates bundling and capping the barb-ends of actin filaments, thereby promoting stress fiber formation and inhibiting actin polymerization which corresponds to our observed changes in actin-binding proteins (Fig. 6; Mitra et al., 2011).

One signaling pathway activated under acute heat stress in *Mytilus* leads to the synthesis of the lipid prostaglandins (PG), lipid messengers which affect the synthesis of stress and inflammation-associated proteins (Duran-Encinas et al., 2024). Central to PG synthesis is the conversion of arachidonic acid to PGH_2_ through cyclcooxygenase (PG-endoperoxide synthase), followed by subsequent conversions to PGE_2_ and PGD_2_ (Fig. 9). LTLF and HTHF mussels increased PGD_2_ isomerase, synthesizing PGD_2_, while LTLF and HTLF increased PG dehydrogenase, which inactivates PGE_2_.

**Figure 9.**
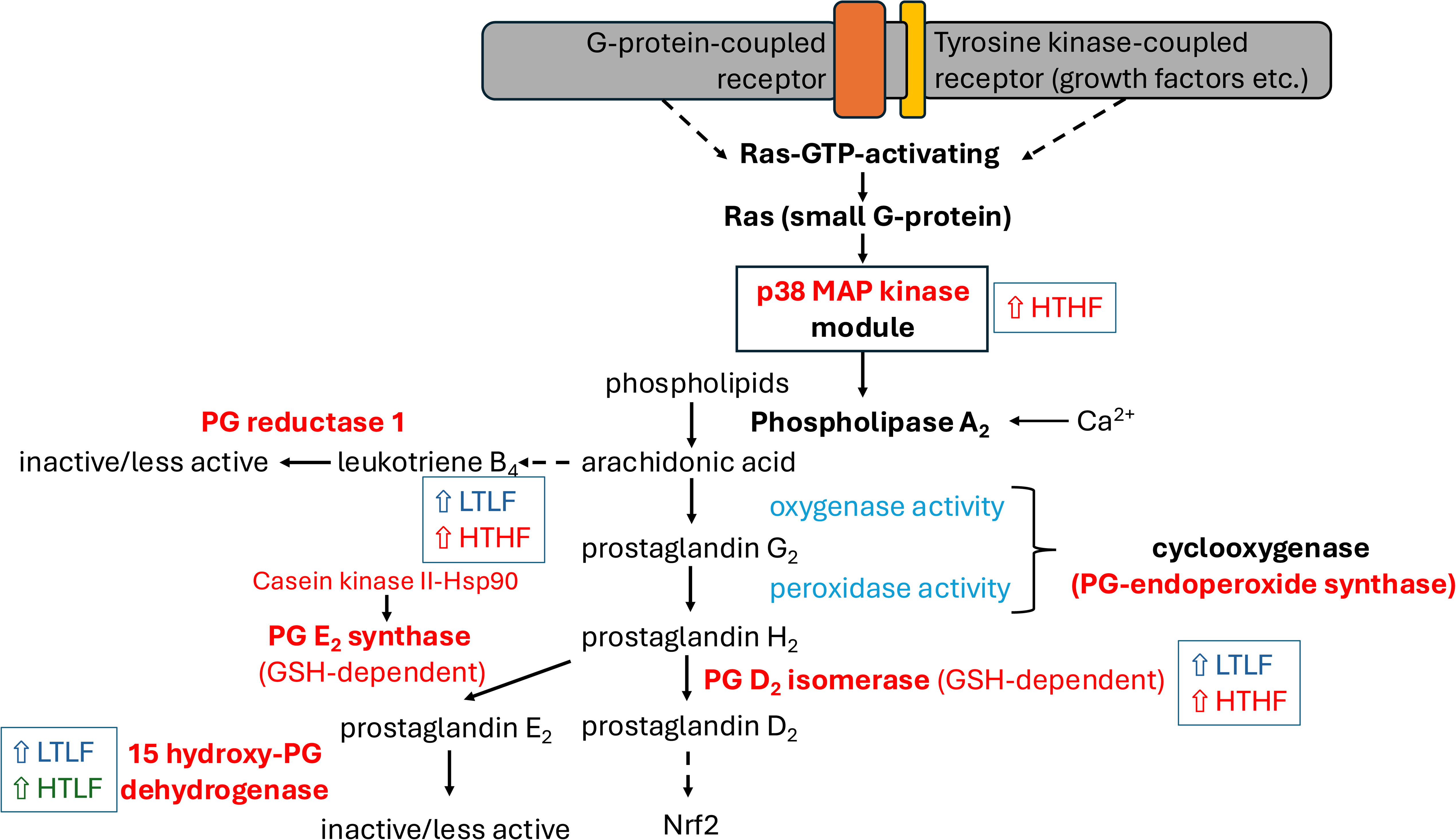
Hypothetical prostaglandin signaling pathway in *Mytilus* gill. The proposed pathway is based on identified signaling proteins leading to the synthesis of prostaglandins D_2_ and E_2_ and indicates the acclimation-dependent changes to acute heat stress.

Other acclimation treatments showed changes in stress-induced signaling proteins. For example, low-temperature-high-food (LTHF) mussels decreased tumor-necrosis factor (TNF) receptor-associated factor 2 (TRAF2), and increased stress-activated protein kinase JNK. TNF controls cell death by apoptosis or necrosis by activating TRAF2 stimulating interactions with kinases such as p38 MAPK, JNK, and the nuclear factor kB (NFkB) modules (Song et al., 1997). The heat-inducible p38 MAPK increased in high-temperature-high-food (HTHF) mussels. Together, this suggests that HF-acclimated mussels increased proteins of two major stress-induced signaling pathways (JNK and p38 MAPK), indicating a robust response to cell stress by activating pro-survival pathways, like the response to acute thermal stress in *Mytilus* species (Anestis et al., 2007; Gourgou et al., 2010). In contrast, the pro-apoptotic tumor protein p53-inducible protein 3, which is activated through DNA damage and regarded as the ‘guardian of the genome’, controls death-associated protein (DAP1) kinase, affects cell adhesion and membrane blebbing and increased in HTHF (Marks et al., 2017). Thus, LTLF mussels changed signaling proteins involved mainly in cytoskeletal and secretory pathway dynamics, while HT-acclimated mussels changed activities of pro-survival and apoptotic pathways.

## DISCUSSION

### mRNA processing and protein synthesis

The proteomic signature of low-temperature-low-food (LTLF) mussels shows a robust increase in mRNA processing proteins, specifically of several splicing factors (SRSF1/9, 4/5/6, 7, and 16) and hnRNPs (A1/A3, F/H, R and U-like 1) throughout recovery. While elements of this response are known to be part of the canonical heat shock response (SRSF 7), others are possibly modulated by the food-dependent remodeling of the translational apparatus, specifically the ribosomal structure, as evidenced by an increase in abundance of structural components of the ribosome in LTLF mussels. This suggests that low food ration requires an immediate restructuring of ribosomes upon heat shock. Interestingly, ribosomal rRNA G+C content correlates with optimal growth temperature in bacteria (Galtier & Lobry, 1997) and likely in many eukaryotes (Varriale et al., 2008), suggesting that rRNA is highly temperature sensitive (Somero et al., 2017). Furthermore, the adjustment of ribosomal structure, involving the interaction between rRNA and ribosomal proteins, to thermal conditions is likely also food dependent, as food ration leads to specialized ribosomes in model eukaryotes (Liu & Qian, 2014; Petibon et al., 2021). Given that the synthesis of paralogous ribosomal proteins and alternative splicing is important for generating ribosomal heterogeneity and modulating translation (Xue & Barna, 2012), it is probable that changes in food ration cause shifts in expression and alternative splicing of ribosomal proteins to adjust mRNA processing to tune translation to prevailing food and thermal conditions.

The sensitivity of the ribosomal composition is illustrated by variation in the response among our acclimation conditions. The composition of ribosomes alters translational control by preferentially selecting specific mRNAs for translation (Shi et al., 2017). Different compositions may also counteract the destabilization of ribosomal subregions during heat stress to preserve translation efficiency (Bell et al., 1988). It appears that this mechanism of selective ribosomal restructuring may be an early response to heat stress to cope with the effects associated with increased temperatures and the sensitivity of ribosomes to oxidative damages (Higgins et al., 2015; Shcherbik & Pestov, 2019; Willi et al., 2018).

Each food-by-temperature condition may require a different strategy in *Mytilus*. While high-temperature-low-food (HTLF) decreased several ribosomal proteins by 25 h, high-temperature-high-food (HTHF) mussels immediately increased several but different ribosomal proteins. These ribosomal restructuring responses of HT-mussels were partially decoupled from mRNA processing, as we observed almost no changes of mRNA processing proteins in these groups. In contrast, low-temperature-high-food (LTHF) mussels showed several changes in mRNA processing, but not as many as LTLF mussels. This suggests that LT-mussels are more food-sensitive in their mRNA processing response than HT-mussels and that HT-mussels are more food-sensitive in remodeling their ribosomes.

Further insight into the differential response comes from a comparison between the RPs changing between LTLF and HTHF mussels. For example, of the three RPs unique to HTHF, the 60S acidic RP P0 interacts with elongation factors by forming a complex with other acidic ribosomal P(1/2) proteins (Tchorzewski, 2002). Furthermore, two unique and one shared RP (L15, S14e, and L11e, respectively) have extra-ribosomal functions as they can activate the tumor suppressor protein p53 through the suppression of its E3 ubiquitin ligase (Bhat et al., 2004).

Together, these changes represent a substantial energy investment but may only partly contribute to greater translation rates. Instead, they may control the mRNA population being translated and activate extra-ribosomal functions as may be the case in HTHF mussels via activation of p53 (Kasteri et al., 2018). It can be assumed that low food (LF-) acclimated mussels have lower protein synthesis rates and lower steady-state levels of ribosomes (Norkko et al., 2005), therefore requiring an increase in synthesis of elements of the translational machinery upon stress. However, the lack of response in HTLF mussels suggests a strong interaction with temperature acclimation, most likely due to a HT-acclimated proteome signature (e.g., oxidative stress proteins), thereby forgoing the need for greater ribosomal protein synthesis or ribosomal restructuring. Restructuring may also enhance selectivity of mRNA of stress-responsive genes, possibly more so in LTLF mussels. Thus, the restructuring of the translation machinery during heat stress is strongly dependent on acclimation conditions, with the most comprehensive changes occurring in LTLF mussels relative to all other mussels.

### Molecular chaperones and cytoskeletal proteins

The dynamics of the actin- and microtubule-associated cytoskeleton are sensitive to stress, including heat and nutrient stress (Williams & Rousseau, 2022). The stability of actin filaments during heat stress is dependent on the action of small Hsps and their phosphorylation (Lavoie et al., 1995). While we found only one small Hsp isoform to change in low temperature-high food (LTHF) mussels, previous proteomic analyses showed several isoforms to be heat-induced and changing with the circadian and circatidal rhythm (Elowe & Tomanek, 2021; Tomanek & Zuzow, 2010). Interestingly, mussels from a similar acclimation experiment but not exposed to acute heat stress (equivalent to our baseline) showed a high diversity of small Hsp isoforms, likely representing different oligomerization states and post-translational modifications (Vasquez et al., in preparation).

Here we identified food-by-temperature-dependent changes for several isoforms of the chaperone-containing tailless complex polypeptide 1 (CCT1) that plays an important role as a chaperone of the actin- and microtubule-associated cytoskeleton (Vallin & Grantham, 2019). For example, increases of several core CCT subunits coincide with the increase in several actin stress fiber stabilizing proteins, suggesting that the complement of CCT subunits stabilizes the cytoskeleton in LTLF mussels and may thereby also reinforce the connection to focal adhesions and the ECM. Exactly how is unclear, but our results suggest that the moonlighting functions of several CCT-subunits fine tune the CSR in a food-by-temperature dependent fashion by supporting actin bundling (CCT-ε) and actin capping (Brackley & Grantham, 2010; Roobol et al., 1999; Svanstrom & Grantham, 2016). The same is likely the case for the interactions between other CCT subunits (CCT-α, −ζ and −η) and the microtubule-associated network. Together, these findings indicate that food-by-temperature conditions set cytoskeletal dynamics and stabilization through molecular chaperone core and moonlighting functions. CCT’s may also trigger the UPR in the ER, at least in starving planarians (Gutierrez-Gutierrez et al., 2021).

### ER chaperoning and secretory pathway

During protein maturation along the ER-Golgi secretory pathways, a variety of oligosaccharides are transferred to produce the O-linked glycans of mucopolysaccharides secreted by gill mucocytes of bivalves which support feeding and pathogen defense (Pales Espinosa et al., 2016). This pathway involves ER chaperones, glucosyl transferases, and glucosidases. While LTLF mussels increased ER chaperoning activity through increases in Grp78, PDIs and PPIs, and the hexosamine pathway (Araki & Nagata, 2012; Braakman & Hebert, 2013), the overall ER and Golgi secretory pathway, including glycosylation, did not (Fig. S1), possibly to save ATP for chaperoning. The consequences for the mucus production that is required for capturing and transporting food particles are unknown and may affect the mucus-defense against pathogens, particularly during stress.

### Stress-induced signaling pathways

Given the many cross-paths and pathway constellations potentially responding to environmental conditions, any signaling pathway assignment based on identification alone is speculative. However, abundance changes and identifications of several canonical members along the prostaglandin synthesis pathway suggest that it is food-by-temperature-dependent. Our results therefore add to the recent confirmation that prostaglandin synthesis changes during acute heat stress (Duran-Encinas et al., 2024).

Prostanoid synthesis is initiated either by G-protein coupled receptors (e.g., neurotransmitter receptors) or by tyrosine (Tyr-)kinase-coupled receptors (e.g., growth factors) and the downstream activation of the small G-protein Ras (Narumiya et al., 1999). Phospholipase A_2_, activated by Ca^2+^ and phosphorylation by MAP kinases, induces the release of arachidonic acid from membrane lipids and thereby triggers the synthesis of eicosanoids, specifically several prostaglandins and leukotrienes (Marks et al., 2017). We identified several Tyr-kinase receptors, associated scaffolding proteins (Grb2), members of Ras family of small G-proteins and p38 MAPK as potential members of a signaling pathway leading to PG synthesis (Figs. 9, S3).

Cyclooxygenase (here PG-endoperoxide synthase) then acts as an oxygenase and peroxidase, leading to prostaglandin H_2_ (PGH_2_), which serves as a precursor for PGE_2_ (PGE_2_ synthase) and PGD_2_ (PGD_2_ isomerase). PGD_2_ isomerase increased in LTLF and HTHF mussels, likely increasing PGD_2_, while 15-hydroxyprostaglandin dehydrogenase increased in in LTLF and HTHF, inactivating PGE_2_ (but not PGD_2_). PGE_2_ itself is produced by PGE_2_ synthase, which is activated through phosphorylation by casein kinase II, and formation to a multi-protein complex with Hsp90, which increased in LTLF and HTHF. And two of these reactions use glutathione as co-factor, coinciding with likely greater glutathione turnover through leucyl aminopeptidase. Thus, these changes suggest that food-by-temperature acclimation effects modify prostaglandin synthesis and are connected to systemic changes.

The roles of prostaglandins in mussels range from osmoregulation and mineral transport, to gametogenesis and immune defense (Rowley et al., 2005; Ruggeri & Thoroughgood, 1985), and have been reported to be synthesized in response to acute rather than chronic heat stress (for PGE_2_ only), coinciding with the recruitment of phagocytes to the gill (Duran-Encinas et al., 2024). Higher PGD_2_ levels in turn can lead to the activation of the antioxidative transcription factor Nrf2 (Ishii, 2015), consistent with increased abundances of several antioxidative stress proteins in this group (Fabela et al., in review). Given that the responses varied among acclimation conditions, PG synthesis patterns warrant deeper studies.

## Conclusion

Our results suggest some novel systemic responses of ciliated epithelial cells and mucocytes, making up most of gill cells, as being fine-tuned by the combination of thermal conditions and the prevailing food ration. The ones presented here are closely linked to changes in redox balance (i.e., oxidative stress) at the cellular level as described in (Fabela et al., in review; May & Tomanek, 2024), and need to be further integrated to tissue and organismal level responses (May et al., 2021). The effect of calorie-dependent NAD-dependent deacylases inhibitors on the proteomic response may give insight into one of the potential regulatory mechanisms (May et al., in preparation). Together, these studies develop a systemic picture of how food availability contributes to determine the cellular stress response.

## Author contributions

Conceptualization: M.A.M., L.T.; Data curation: F.R.F., M.A.M.; Formal analysis: F.R.F., M.A.M., L.T.; Funding acquisition: F.R.F., L.T.; Investigation: F.R.F., M.A.M., L.T.; Methodology: F.R.F., M.A.M., L.T.; Project administration: M.A.M., L.T.; Resources: L. T.; Supervision: M.A.M., L.T.; Writing – original draft: L.T., M.A.M., F.R.F.; Writing – review & editing: F.R.F., M.A.M., L.T.

## ACKNOWLEDGMENTS

The authors are grateful for the assistance of numerous undergraduate and graduate students in conducting these experiments, with special thanks to Drs. M. Christina Vasquez and Anne Todgham. The success of these projects would not have been possible without the logistical support from the Cal Poly Biology Department and Center for Coastal Marine Studies, including Kevin Dunham, Rob Brewster, Doug Brewster, and Jason Felton.

## DATA AND RESOURCE AVAILABILITY

The mass spectrometry proteomics data have been deposited to the ProteomeXchange Consortium (http://proteomecentral.proteomexchange.org) via the PRoteomics IDentifications Database (PRIDE) (Perez-Riverol et al., 2025) partner repository with the dataset identifier PXD072074 and 10.6019/PXD072074. The experimental data are deposited on Dryad under DOI: 10.5061/dryad.pc866t24k.

## SUPPLEMENT

**Figure S1.**
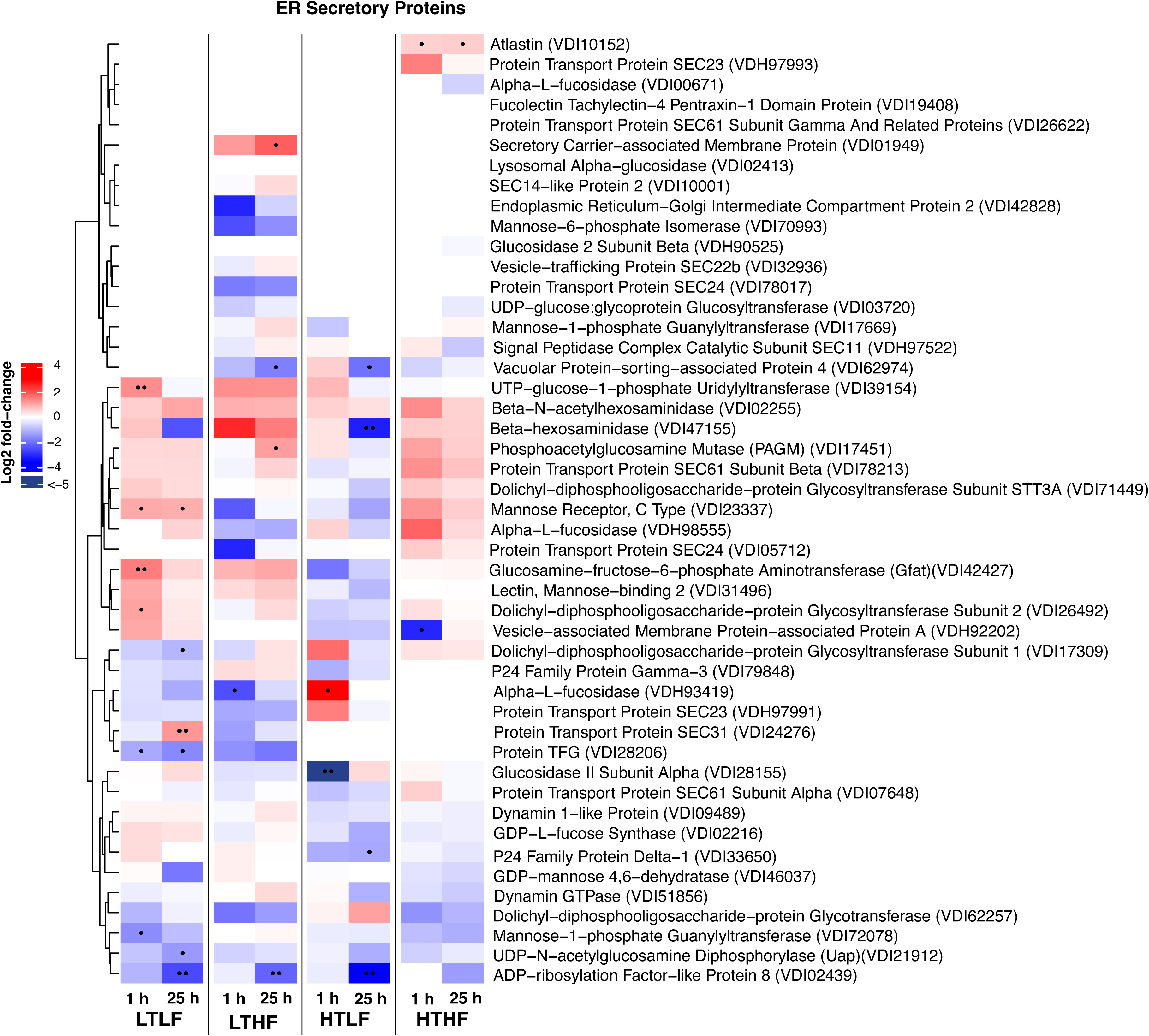
Heat map of ER Golgi secretory pathway proteins. The heat map shows log_2_ fold change in protein abundance from 1 h or 25 h recovery relative to the baseline in each of the acclimation groups.: low temperature-low food (LTLF), low temperature-high food (LTHF), high temperature-low food (HTLF), and high temperature-high food (HTHF). Increases in abundance are indicated by warm colors (reds), while decreases are cool (blues). Significance levels are shown by dots, where ••• signifies p < 0.001, •• is p < 0.01, and • is p < 0.05. Proteins were identified using GenBank accession numbers.

**Figure S2.**
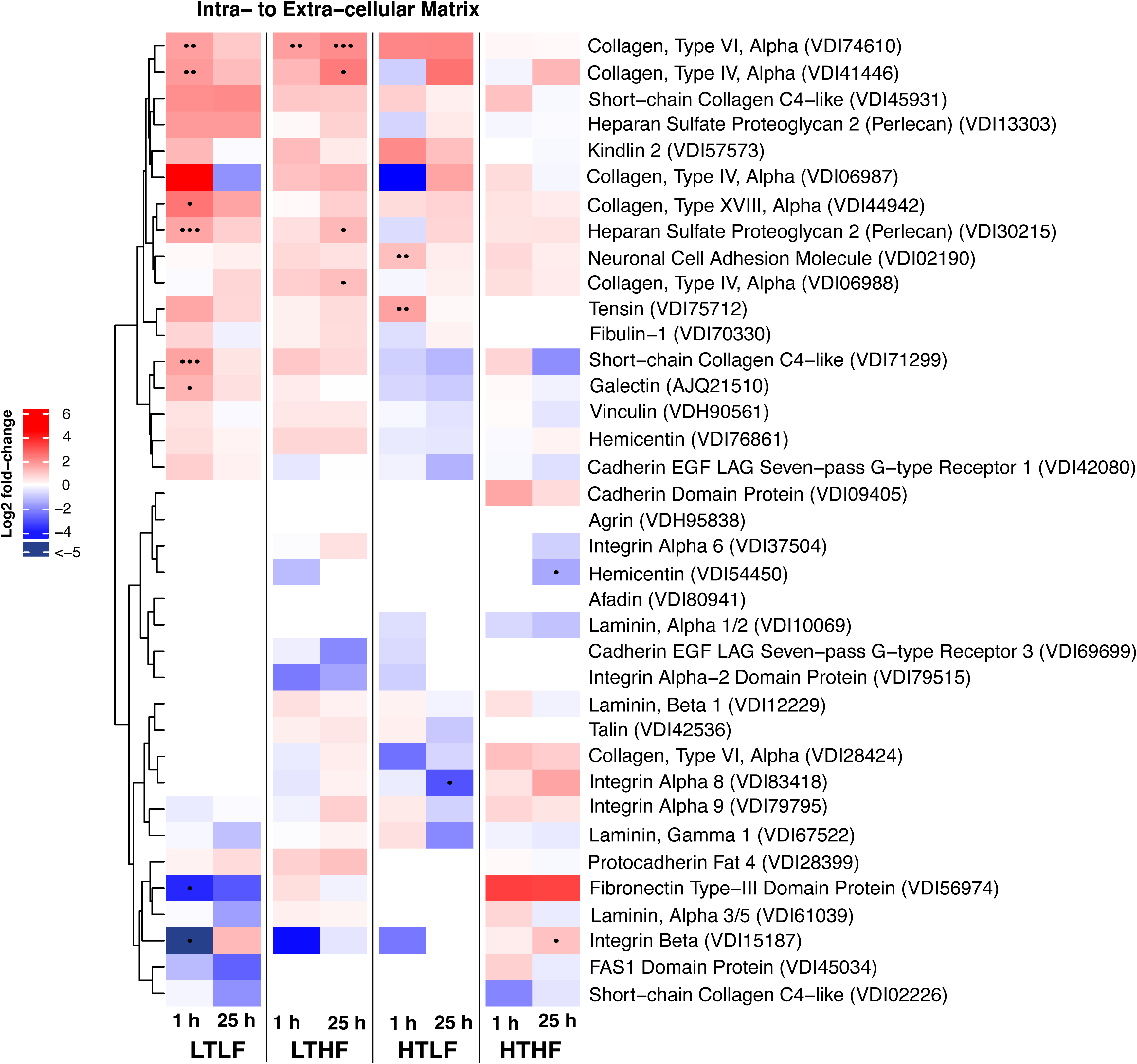
Heat map of intracellular to extra-cellular matrix proteins. The heat map shows log_2_ fold change in protein abundance from 1 h or 25 h recovery relative to the baseline in each of the acclimation groups; For additional details see Fig. S1.

**Figure S3.**
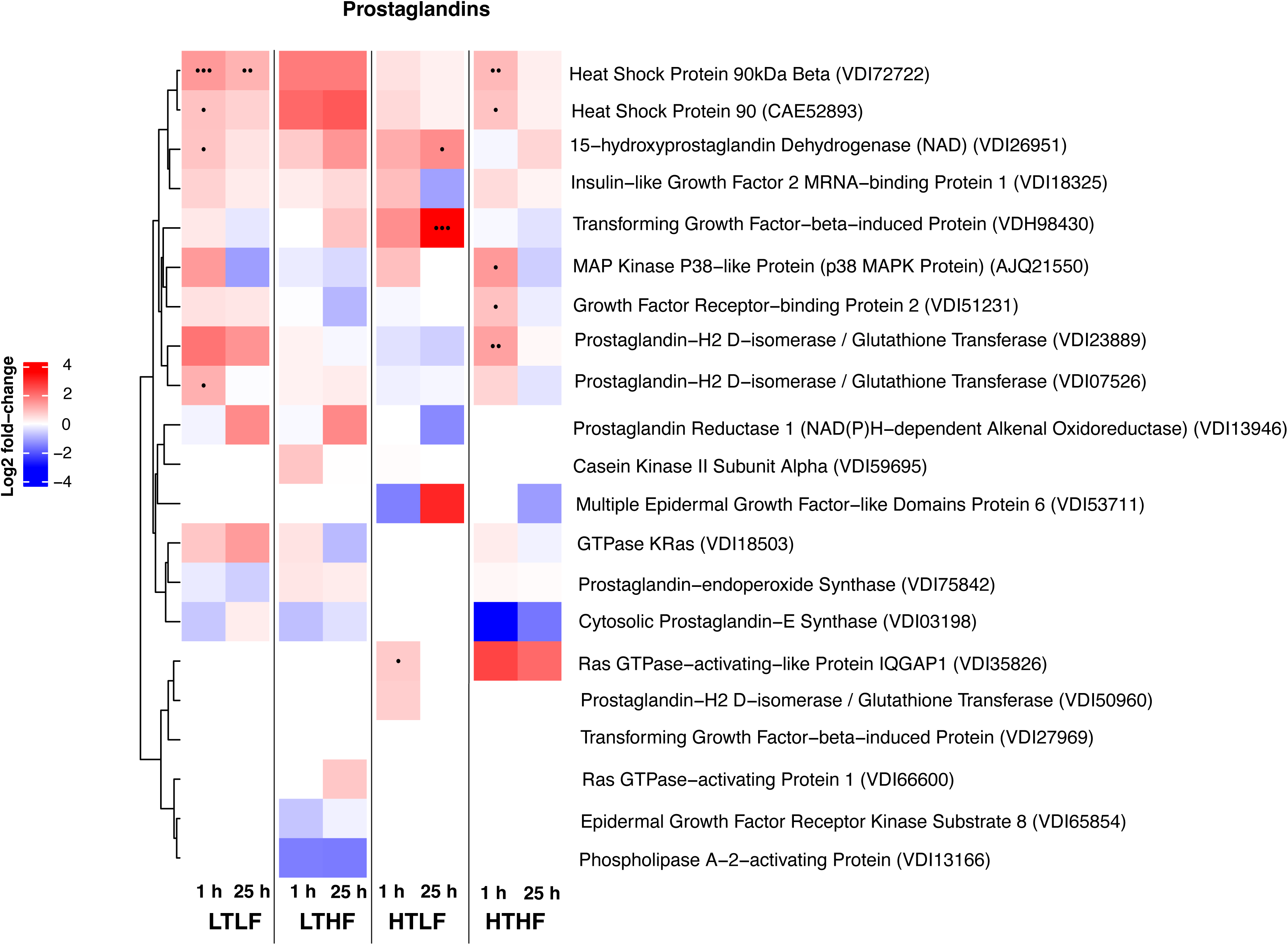
Heat map of hypothetical prostaglandin (PG) synthesis pathway proteins. The heat map shows log_2_ fold change in protein abundance from 1 h or 25 h recovery relative to the baseline in each of the acclimation groups; For additional details see Fig. S3.

