## Supplemental Material for "Food Ration Affects mRNA Processing, Translation, Proteostasis, and Cytoskeletal Responses During Heat Shock in *Mytilus californianus*"

06/02/2026

### SUPPLEMENT

Tomanek[1], L., Fabela[1], R. F., and May[1, 2], M. A.

[1]: California Polytechnic State University, Department of Biological Sciences, 1 Grand Ave., San Luis Obispo, CA 93407, USA.

[2]: Florida Gulf Coast University, Department of Marine and Earth Sciences, 10501 FGCU Blvd. S, Fort Myers, FL  33965, USA

**Figure S1. Heat map of ER Golgi secretory pathway proteins.** The heat map shows $log_2$ fold change in protein abundance from 1 h or 25 h recovery relative to the baseline in each of the acclimation groups.: low temperature-low food (LTLF), low temperature-high food (LTHF), high temperature-low food (HTLF), and high temperature-high food (HTHF). Increases in abundance are indicated by warm colors (reds), while decreases are cool (blues). Significance levels are shown by dots, where ••• signifies $p < 0.001$, •• is $p < 0.01$, and • is $p < 0.05$. Proteins were identified using GenBank accession numbers.

**Figure S2. Heat map of intracellular to extra-cellular matrix proteins.** The heat map shows $log_2$ fold change in protein abundance from 1 h or 25 h recovery relative to the baseline in each of the acclimation groups; For additional details see Fig. S1.

**Figure S3. Heat map of hypothetical prostaglandin (PG) synthesis pathway proteins.** The heat map shows $log_2$ fold change in protein abundance from 1 h or 25 h recovery relative to the baseline in each of the acclimation groups; For additional details see Fig. S3.

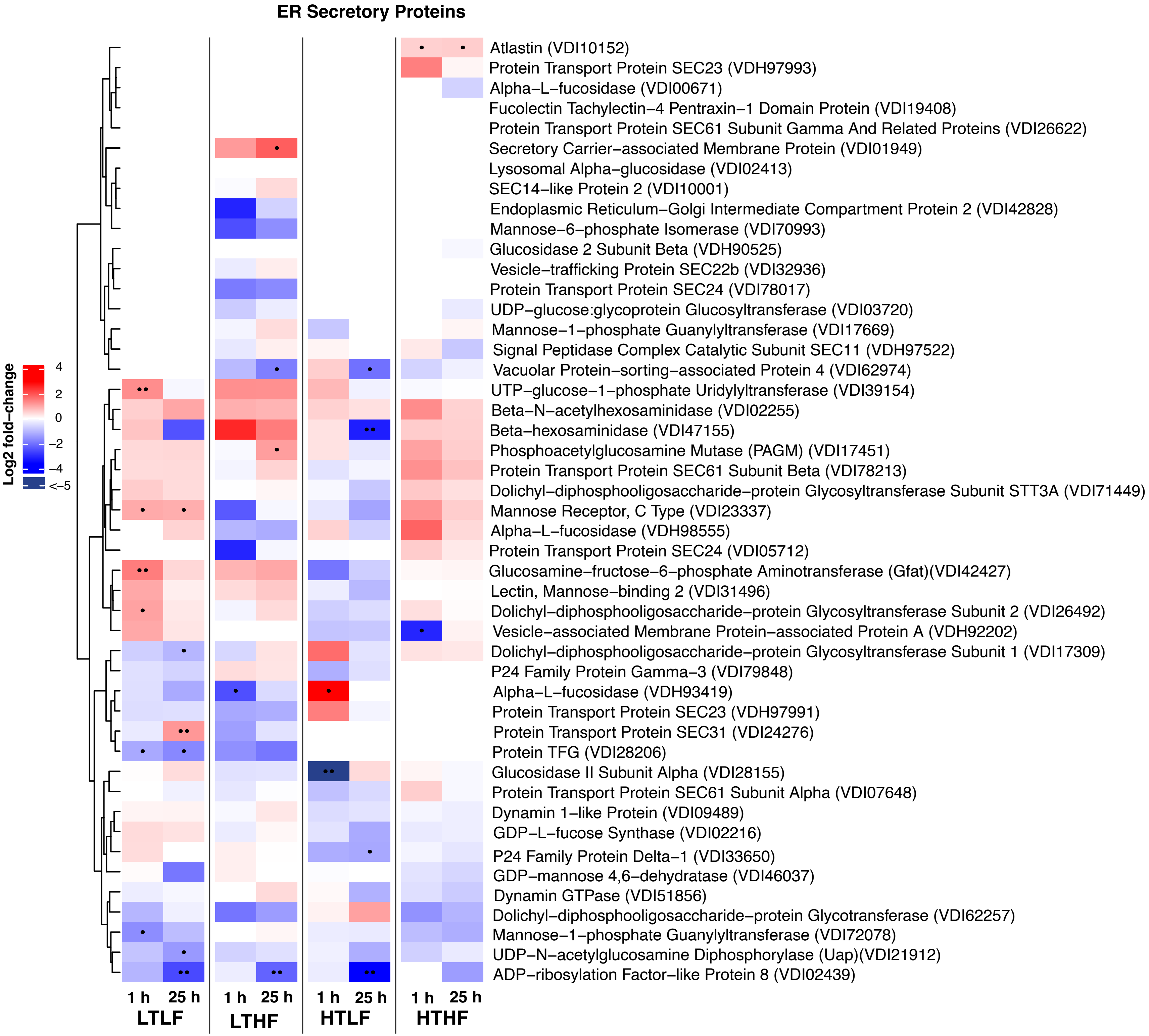
ER Secretory Proteins
Log2 fold−change
4
2
0
−2
−4
<−5
Atlastin (VDI10152)
Protein Transport Protein SEC23 (VDH97993)
Alpha−L−fucosidase (VDI00671)
Fucolectin Tachylectin−4 Pentraxin−1 Domain Protein (VDI19408)
Protein Transport Protein SEC61 Subunit Gamma And Related Proteins (VDI26622)
Secretory Carrier−associated Membrane Protein (VDI01949)
Lysosomal Alpha−glucosidase (VDI02413)
SEC14−like Protein 2 (VDI10001)
Endoplasmic Reticulum−Golgi Intermediate Compartment Protein 2 (VDI42828)
Mannose−6−phosphate Isomerase (VDI70993)
Glucosidase 2 Subunit Beta (VDH90525)
Vesicle−trafficking Protein SEC22b (VDI32936)
Protein Transport Protein SEC24 (VDI78017)
UDP−glucose:glycoprotein Glucosyltransferase (VDI03720)
Mannose−1−phosphate Guanylyltransferase (VDI17669)
Signal Peptidase Complex Catalytic Subunit SEC11 (VDH97522)
Vacuolar Protein−sorting−associated Protein 4 (VDI62974)
UTP−glucose−1−phosphate Uridylyltransferase (VDI39154)
Beta−N−acetylhexosaminidase (VDI02255)
Beta−hexosaminidase (VDI47155)
Phosphoacetylglucosamine Mutase (PAGM) (VDI17451)
Protein Transport Protein SEC61 Subunit Beta (VDI78213)
Dolichyl−diphosphooligosaccharide−protein Glycosyltransferase Subunit STT3A (VDI71449)
Mannose Receptor, C Type (VDI23337)
Alpha−L−fucosidase (VDH98555)
Protein Transport Protein SEC24 (VDI05712)
Glucosamine−fructose−6−phosphate Aminotransferase (Gfat)(VDI42427)
Lectin, Mannose−binding 2 (VDI31496)
Dolichyl−diphosphooligosaccharide−protein Glycosyltransferase Subunit 2 (VDI26492)
Vesicle−associated Membrane Protein−associated Protein A (VDH92202)
Dolichyl−diphosphooligosaccharide−protein Glycosyltransferase Subunit 1 (VDI17309)
P24 Family Protein Gamma−3 (VDI79848)
Alpha−L−fucosidase (VDH93419)
Protein Transport Protein SEC23 (VDH97991)
Protein Transport Protein SEC31 (VDI24276)
Protein TFG (VDI28206)
Glucosidase II Subunit Alpha (VDI28155)
Protein Transport Protein SEC61 Subunit Alpha (VDI07648)
Dynamin 1−like Protein (VDI09489)
GDP−L−fucose Synthase (VDI02216)
P24 Family Protein Delta−1 (VDI33650)
GDP−mannose 4,6−dehydratase (VDI46037)
Dynamin GTPase (VDI51856)
Dolichyl−diphosphooligosaccharide−protein Glycotransferase (VDI62257)
Mannose−1−phosphate Guanylyltransferase (VDI72078)
UDP−N−acetylglucosamine Diphosphorylase (Uap)(VDI21912)
ADP−ribosylation Factor−like Protein 8 (VDI02439)
1 h
25 h
LTLF
1 h
25 h
LTHF
1 h
25 h
HTLF
1 h
25 h
HTHF

Fig. S2

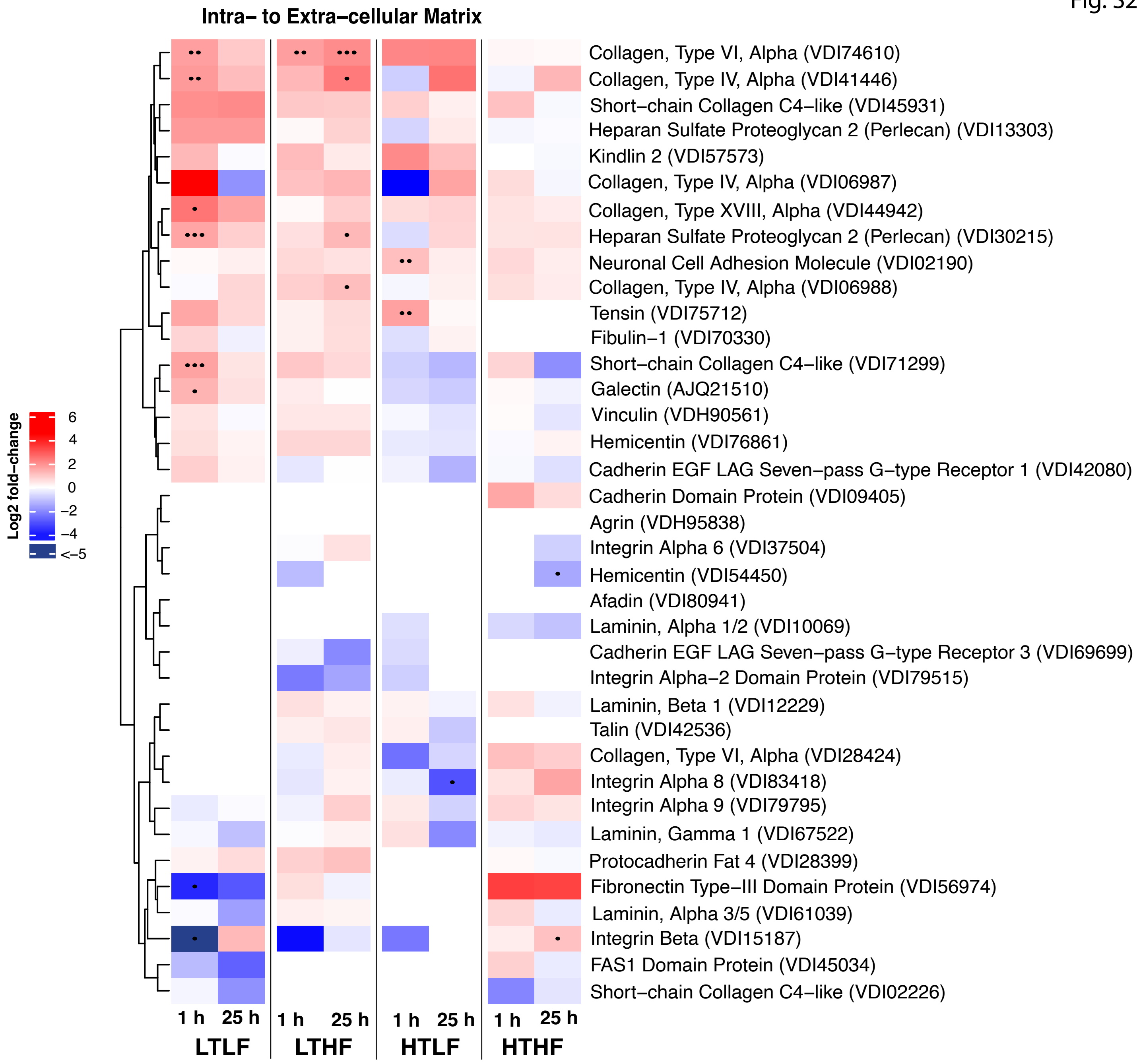
Intra– to Extra–cellular Matrix
Log2 fold–change
6
4
2
0
–2
–4
<–5
Collagen, Type VI, Alpha (VDI74610)
Collagen, Type IV, Alpha (VDI41446)
Short–chain Collagen C4–like (VDI45931)
Heparan Sulfate Proteoglycan 2 (Perlecan) (VDI13303)
Kindlin 2 (VDI57573)
Collagen, Type IV, Alpha (VDI06987)
Collagen, Type XVIII, Alpha (VDI44942)
Heparan Sulfate Proteoglycan 2 (Perlecan) (VDI30215)
Neuronal Cell Adhesion Molecule (VDI02190)
Collagen, Type IV, Alpha (VDI06988)
Tensin (VDI75712)
Fibulin–1 (VDI70330)
Short–chain Collagen C4–like (VDI71299)
Galectin (AJQ21510)
Vinculin (VDH90561)
Hemicentin (VDI76861)
Cadherin EGF LAG Seven–pass G–type Receptor 1 (VDI42080)
Cadherin Domain Protein (VDI09405)
Agrin (VDH95838)
Integrin Alpha 6 (VDI37504)
Hemicentin (VDI54450)
Afadin (VDI80941)
Laminin, Alpha 1/2 (VDI10069)
Cadherin EGF LAG Seven–pass G–type Receptor 3 (VDI69699)
Integrin Alpha–2 Domain Protein (VDI79515)
Laminin, Beta 1 (VDI12229)
Talin (VDI42536)
Collagen, Type VI, Alpha (VDI28424)
Integrin Alpha 8 (VDI83418)
Integrin Alpha 9 (VDI79795)
Laminin, Gamma 1 (VDI67522)
Protocadherin Fat 4 (VDI28399)
Fibronectin Type–III Domain Protein (VDI56974)
Laminin, Alpha 3/5 (VDI61039)
Integrin Beta (VDI15187)
FAS1 Domain Protein (VDI45034)
Short–chain Collagen C4–like (VDI02226)
1 h
25 h
LTLF
1 h
25 h
LTHF
1 h
25 h
HTLF
1 h
25 h
HTHF

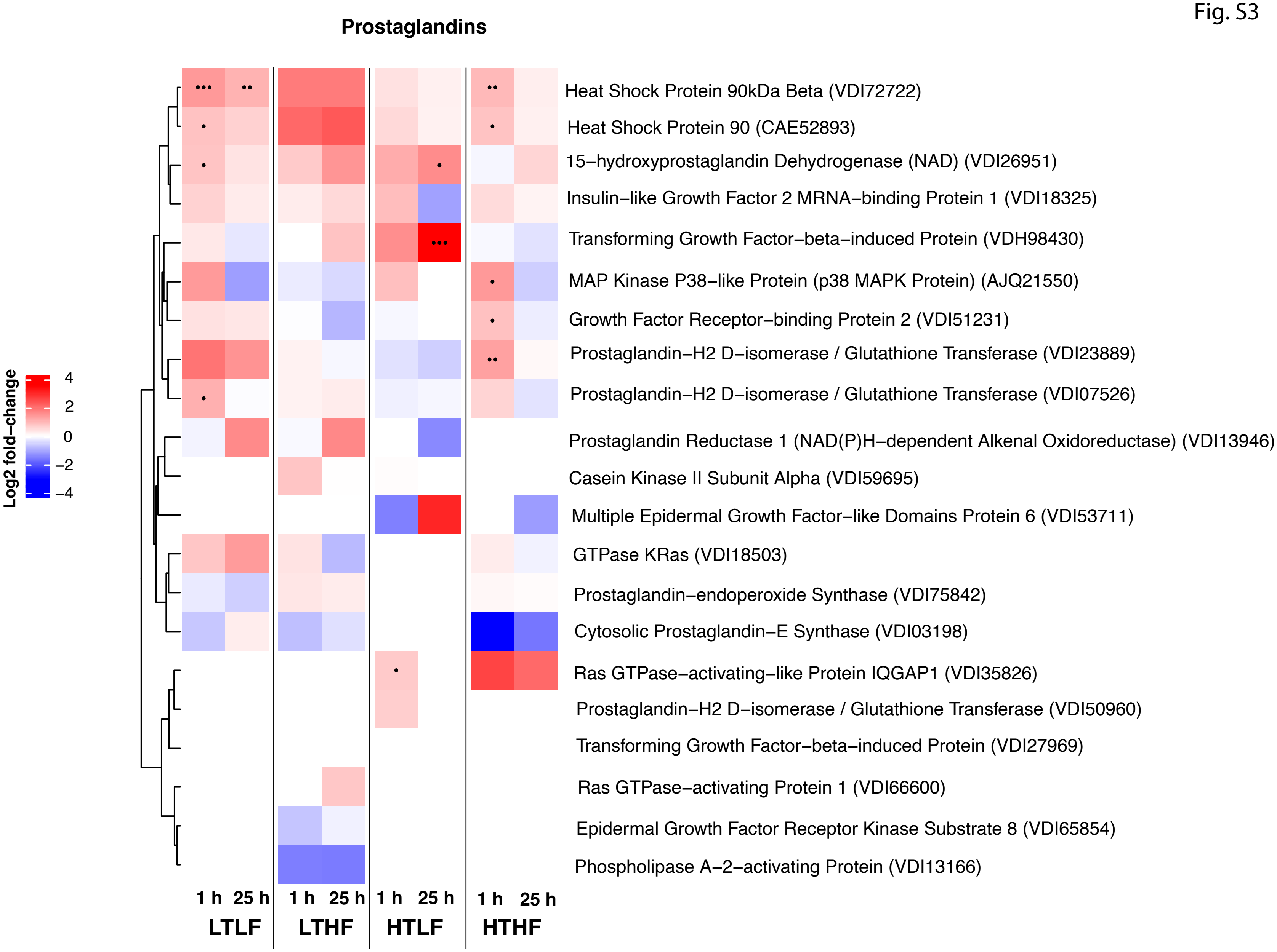
Prostaglandins
Log2 fold−change
4
2
0
−2
−4
Heat Shock Protein 90kDa Beta (VDI72722)
Heat Shock Protein 90 (CAE52893)
15−hydroxyprostaglandin Dehydrogenase (NAD) (VDI26951)
Insulin−like Growth Factor 2 MRNA−binding Protein 1 (VDI18325)
Transforming Growth Factor−beta−induced Protein (VDH98430)
MAP Kinase P38−like Protein (p38 MAPK Protein) (AJQ21550)
Growth Factor Receptor−binding Protein 2 (VDI51231)
Prostaglandin−H2 D−isomerase / Glutathione Transferase (VDI23889)
Prostaglandin−H2 D−isomerase / Glutathione Transferase (VDI07526)
Prostaglandin Reductase 1 (NAD(P)H−dependent Alkenal Oxidoreductase) (VDI13946)
Casein Kinase II Subunit Alpha (VDI59695)
Multiple Epidermal Growth Factor−like Domains Protein 6 (VDI53711)
GTPase KRas (VDI18503)
Prostaglandin−endoperoxide Synthase (VDI75842)
Cytosolic Prostaglandin−E Synthase (VDI03198)
Ras GTPase−activating−like Protein IQGAP1 (VDI35826)
Prostaglandin−H2 D−isomerase / Glutathione Transferase (VDI50960)
Transforming Growth Factor−beta−induced Protein (VDI27969)
Ras GTPase−activating Protein 1 (VDI66600)
Epidermal Growth Factor Receptor Kinase Substrate 8 (VDI65854)
Phospholipase A−2−activating Protein (VDI13166)
1 h 25 h
1 h 25 h
1 h 25 h
1 h 25 h
LTLF
LTHF
HTLF
HTHF
